# *SFPQ-TFE3* gene fusion reciprocally regulates mTORC1 activity and induces lineage plasticity in a novel mouse model of renal tumorigenesis

**DOI:** 10.1101/2024.11.21.624702

**Authors:** Kaushal Asrani, Adrianna Amaral, Juhyung Woo, Sanaz Nourmohammadi Abadchi, Thiago Vidotto, Kewen Feng, Hans B. Liu, Mithila Kasbe, Masaya Baba, Yuichi Oike, Patricia Outeda, Terry Watnick, Avi Z. Rosenberg, Laura S. Schmidt, W. Marston Linehan, Pedram Argani, Tamara L. Lotan

**Author notes:** **Correspondence**: Tamara Lotan, MD, 1550 Orleans Street, Baltimore, MD 21231. Lead Contact Correspondence (KA), (TLL). Equal contribution (Kaushal Asrani, Adrianna Amaral).

## Abstract

The MiT/TFE family gene fusion proteins, such as *SFPQ-TFE3*, drive both epithelial (eg, translocation renal cell carcinoma, tRCC) and mesenchymal (eg, perivascular epithelioid cell tumor, PEComa) neoplasms with aggressive behavior. However, no prior mouse models for *SFPQ-TFE3*-related tumors exist and the mechanisms of lineage plasticity induced by this fusion remain unclear. Here, we demonstrate that constitutive murine renal expression of human *SFPQ-TFE3* using Ksp Cadherin-Cre as a driver disrupts kidney development leading to early neonatal renal failure and death. In contrast, post-natal induction of *SFPQ-TFE3* in renal tubular epithelial cells using Pax8 ERT-Cre induces infiltrative epithelioid tumors, which morphologically and transcriptionally resemble human PEComas. As seen in MiT/TFE fusion-driven human tumors, *SFPQ-TFE3* expression is accompanied by the strong induction of mTORC1 signaling, which is partially amino acid-sensitive and dependent on increased *SFPQ-TFE3*-mediated *RRAGC/D* transcription. Remarkably, *SFPQ-TFE3* expression is sufficient to induce lineage plasticity in renal tubular epithelial cells, with rapid down-regulation of the critical PAX2/PAX8 nephric lineage factors and tubular epithelial markers, and concomitant up-regulation of PEComa differentiation markers in transgenic mice, human cell line models and human tRCC. Pharmacologic or genetic inhibition of mTOR signaling downregulates expression of the *SFPQ-TFE3* fusion protein and rescues nephric lineage marker expression and transcriptional activity *in vitro.* These data provide evidence of a potential epithelial cell-of-origin for *TFE3*-driven PEComas and highlight a reciprocal role for *SFPQ-TFE3* and mTOR in driving lineage plasticity in the kidney, expanding our understanding of the pathogenesis of MiT/TFE-driven tumors.

## Introduction

Translocation renal cell carcinoma (tRCC) is a rare subtype of non-clear cell, sporadic kidney cancer driven by chromosomal translocations involving the MiT/TFE family of transcription factors (*TFE3* [Xp11.23], *TFEB* [6p21.1], and *MITF* [3p13]), which are key regulators of lysosomal biogenesis. The resulting fusions of MiT/TFE genes with various partner genes (the most common being *ASPSCR1, PRCC,* and *SFPQ*), leads to constitutive nuclear localization and activation of the chimeric transcription factors^1,2^. tRCC exhibit a high degree of morphologic and clinical heterogeneity, at least in part due to varying *TFE3* fusion partners^2^, and their molecular landscape has only been partially defined, prompting an urgent need to identify novel biomarkers and therapeutic targets.

In addition to renal carcinomas, *TFE3*-fusions are also implicated in the pathogenesis of a subset of potentially aggressive mesenchymal tumors called perivascular epithelioid cell tumors (PEComas), which are derived from an unknown cell-of-origin, occurring in the kidney and many other sites. tRCC and PEComas have highly overlapping epithelioid morphologies and immunophenotypes, including expression of melanocytic lineage genes and lysosomal markers (PMEL, MelanA, Cathepsin K, GPNMB) that are canonical MiT/TFE transcriptional targets^3–5^. Pathologically, the key distinction between tRCC and MiT/TFE-driven renal PEComas is the lack of apparent epithelial differentiation in the latter, as evidenced by the complete absence of pan-epithelial markers (cytokeratins/EMA/CD10)^6^ and loss of critical renal lineage transcription factors, PAX8 and PAX2^7–9^. In contrast, there is retention of these renal epithelial lineage markers in tRCC, similar to what is seen in clear cell RCC, where PAX8 is essential for oncogenesis^10,11^.

Though some have posited that PEComas originate from pericytes^12^ or neural crest cells^13,14^, at least one study has indicated a potential clonal origin of renal PEComa cells from tumor-initiating, renal proximal tubular epithelial cells ^15^ and hybrid epithelial/mesenchymal tumors have been described in patients^16^. This fact, together with the frequent underexpression of cytokeratin and EMA expression seen in tRCC cases^17^, suggests the possibility that tRCC and renal PEComas may develop from the same epithelial cell-of-origin, which subsequently undergoes transdifferentiation in the latter. However no *in vivo* transgenic models of PEComa tumorigenesis have been described to date, hampering progress in understanding mechanisms of tumorigenesis and potential therapies for this rare, but potentially aggressive, tumor type.

Though rearrangements involving the MiT/TFE genes *TFE3* and *TFEB* are critical drivers for a subset of PEComas, a majority of renal PEComas (including the most common and well-differentiated subtype known as angiomyolipomas) are driven by alternate genomic alterations, most commonly biallelic *TSC1/2* loss resulting in mTORC1 hyperactivation. Explaining the mutual exclusivity between *TSC1/2* inactivation and MiT/TFE gene rearrangements in PEComas, we and others have previously shown that mTORC1 activation leads to constitutive activation of *TFEB* and *TFE3* through *FLCN* inactivation^18,19^. Accordingly, inactivating alterations in *FLCN* also drive a rare subset of PEComas, in addition to renal carcinomas that bear resemblance to tRCC ^20–22^. Of note, the mTOR pathway is itself strongly activated in tRCC via largely undefined mechanisms; multiple studies have shown enrichment of mTOR signaling in human tRCC^17,23–25^, as well as cell-line^26^, PDX^27^, and one transgenic mouse tRCC model^25^. Recent studies in renal carcinoma systems have underscored the reciprocity of this mTORC1-MiT interaction in the kidney: in the setting of *TSC1/2* or *FLCN* loss, TFEB and/or TFE3 are constitutively activated, drive renal tumorigenesis and these transcription factors are also upstream drivers of mTOR signaling ^18,19,28,29^, indicating that the persistent co-activation of catabolic (autophagy and lysosomal biogenesis) and anabolic (mTORC1) transcriptional programs is a hallmark - albeit paradoxical - feature of many types of renal tumors.

Here, we describe a novel transgenic mouse model of renal tumorigenesis induced by *SFPQ-TFE3* expression, the most common *TFE3* gene fusion seen in human PEComas and their melanotic variants^9,30,31^, and the last of the three most common tRCC fusions that has yet to be modeled in mice. We demonstrate that *SFPQ-TFE3* induces the development of renal tumors recapitulating human PEComas when constitutively expressed in postnatal renal tubular epithelial cells, constituting the first transgenic model system for this rare tumor type. Remarkably, *SFPQ-TFE3* expression is sufficient to rapidly drive lineage plasticity in renal tubular cells, evidenced by PAX2/8 downregulation, loss of cytokeratin expression and melanocytic/lysosomal marker upregulation. MiT/TFE fusion-driven renal tumorigenesis is accompanied by early activation of the mTORC1 pathway across multiple model systems, due at least in part to increased MiT/TFE-mediated *RRAGC/D* transcription. Highlighting the persistently bi-directional regulation of MiT/TFE activity and mTOR, mTOR signaling inhibition is sufficient to rescue renal lineage marker expression and transcriptional activity by downregulating *TFE3*-fusion expression. Taken together, these transgenic models highlight a novel form of epithelial lineage plasticity driven by *TFE3*-fusions and implicate the critical role of mTOR signaling in MiT/TFE fusion-driven tumor types.

## Results

### Generation of an inducible murine allele of SFPQ-TFE3

We generated a transgenic mouse which conditionally expresses the human *SFPQ-TFE3* fusion, downstream of a *LoxP-Stop-LoxP (LSL)* cassette, under control of a strong chicken beta-actin (CAG) promoter in the Rosa 26 locus (*SFPQ-TFE3^LSL^;* hereafter referred to as *ST* mice). Briefly, the *LSL-SFPQ-TFE3* allele, containing the human *SFPQ* CDS (exon 1-9) /human *TFE3* CDS (exon 5-10) (type 2 fusion)^1^, was cloned into intron 1 of the *Rosa26* locus in reverse orientation, separated from the native CAG promoter by a stop sequence flanked by *LoxP* sites *(LSL)*, thus enabling conditional excision of the stop sequence upon Cre-mediated recombination and expression of the fusion transgene (**Fig. S1A**). To validate conditional expression of the allele, renal tubular epithelial cells from adult C57BL/6 controls or *ST* transgenic mice were harvested, dissociated, and cultured in the presence of a control adenovirus or adenovirus expressing Cre-recombinase. Primary cells from *ST*, but not control mice, showed strong induction of SFPQ-TFE3 expression and the canonical MiT/TFE E-box target GPNMB^3^ on immunoblotting (**Fig. S1B)**. Genotypes were additionally confirmed by PCR (**Fig. S1C, D)**.

### Ksp-Cadherin Cre-mediated induction of SFPQ-TFE3 disrupts kidney development with renal failure and early neonatal death

To conditionally express *SFPQ-TFE3* in tubular epithelial cells during renal development, we crossed *ST* transgenic mice with Ksp-Cadherin (Cadherin-16 [Cdh16]) -Cre mice, in which the Cre recombinase is specifically expressed from the Cdh16 promoter in distal tubular epithelial cells and collecting ducts of the kidney starting from embryonic day E12.5^32^, to generate *SFPQ-TFE3^LSL^; Ksp-Cre* mice (hereafter referred to as *STK* mice). While the *STK* mice were born in expected mendelian ratios and did not manifest any gross anatomic differences compared to littermate controls at birth, no *STK* mice survived beyond postnatal day 20 (P20) (**Fig. 1A**). Histologically, *STK* kidneys had a disorganized appearance at P1 with mild and scattered tubular dilation and abundant expression of nuclear TFE3 in tubular epithelial cells by IHC (**Fig. 1B**). By P15, *STK* kidneys were grossly enlarged and markedly cystic, with disruption of the cortical architecture (**Fig. 1C, D**), and significantly elevated kidney to body weight ratios, as well as blood urea nitrogen (BUN) and serum creatinine (**Fig. E-G**). Indirect immunofluorescence for LTL, which labels the brush border of differentiated proximal tubules, showed diminished staining in the *STK* kidney tubules, compared to littermate controls, with rare dilated tubules expressing distal tubular markers such as DBA (**Fig. 1H**). Histologic analysis of kidneys from 3-week-old *STK* mice demonstrated tubular enlargement by solid nests of large epithelioid cells with abundant clear-to-eosinophilic cytoplasm and monomorphic nuclei (**Fig. 1I**). The mutant kidneys also showed luminal occlusion with frequent psammomatous calcification (**Fig. 1I**), a feature of human tRCC^5^, and trichrome staining highlighted inter-tubular fibrosis (**Fig. S2A)**. By P15, there was diffuse tubular nuclear TFE3 expression in *STK* kidneys, and upregulation of an array of melanocytic/lysosomal markers over-expressed in human MiT/TFE-related neoplasms (PMEL, MelanA and GPNMB)^3,5,30^ (**Fig. 1J-L**). Consistent with a pro-tumorigenic role of SFPQ-TFE3, proliferation as measured by BrDU positivity, Ki67 staining and phosphorylated H3 were all significantly elevated in *STK* kidneys compared to littermate controls by 2 weeks of age (**Fig. S2B-D)**. Cumulatively, these results indicate that neonatal expression of renal *SFPQ-TFE3* disrupts renal development with subsequent renal failure culminating in early postnatal death. Though there was evidence of increased tubular cell proliferation, the early renal failure precluded aging the mice to assess for tumor development in *STK* model.

**Figure 1:**
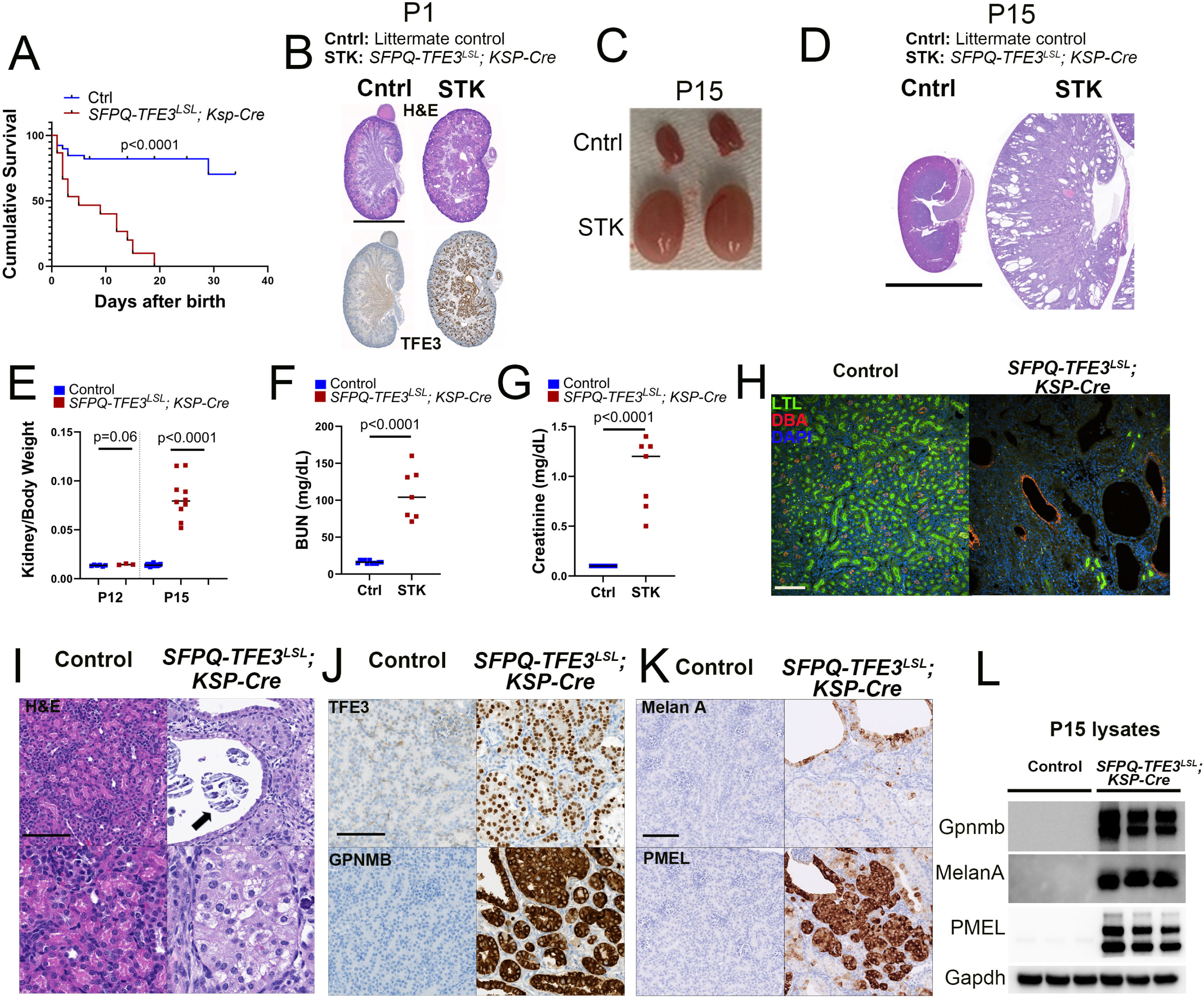
Ksp-Cadherin Cre-mediated induction of SFPQ-TFE3 disrupts kidney development with renal failure and early neonatal death. **(A)** Kaplan-Meier survival analyses of age-matched, control and *SFPQ-TFE3^LSL^; Ksp-Cre* transgenic mice. The following numbers were analyzed: [control=39, STK=15]. P<0.0001 by Log-rank (Mantel-Cox) test. **(B)** Hematoxylin and eosin (H&E) and TFE3 IHC staining of kidneys from age-matched (1 day old), control and *SFPQ-TFE3^LSL^; Ksp-Cre* transgenic mice. Scale bar: 1 mm. **(C)** Macroscopic images of kidneys from age-matched (15 days old), control and *SFPQ-TFE3^LSL^; Ksp-Cre* transgenic mice. **(D)** H&E images of kidneys from age-matched (15 days old), control and *SFPQ-TFE3^LSL^; Ksp-Cre* transgenic mice. Scale bar: 5 mm. **(E)** Kidney to body weight ratios of age-matched, control and *SFPQ-TFE3^LSL^; Ksp-Cre* transgenic mice on post-natal day 12 and day 15. The following numbers were analyzed: a) Kidneys to body weight ratios at day 12 [control=6, STK=3] and day 15 [control=17 and STK=10]. P values by Mann-Whitney test. (**F)** BUN levels and (**G)** serum creatinine levels of the indicated genotypes of mice at day 15. The following numbers were analyzed: [control=11, STK=7]. P values by Mann-Whitney test. **(H)** Indirect immunofluorescence for LTL (green) and DBA (red) in the indicated genotypes of mice on post-natal day 15. Scale bar=100 µm. (**I)** Low (top) and high (bottom) magnification images of H&E-stained kidneys of the indicated genotypes. Arrow indicates intratubular psammomatous calcification. Scale bar=100 µm. Representative immunohistochemistry (IHC) for (**J)** TFE3 (top row), GPNMB (bottom row) **(K)** Melan A (top row), and PMEL (bottom row) in the indicated genotypes of mice on post-natal day 15. Scale bar= 100 µm. **(L)** Immunoblotting of kidney lysates from control and *SFPQ-TFE3^LSL^; Ksp-Cre* transgenic mice at post-natal day 15 for the indicated antibodies. Data are presented as median. Source data are provided as a Source data file.

### Conditional post-natal induction of SFPQ-TFE3 in Pax8 Cre-ER^T2^ mice induces renal tumor development

To examine effects of SFPQ-TFE3 expression in renal tubular cells following completion of kidney development, we leveraged a conditional, tamoxifen-inducible Pax8 Cre-ER^T2^ model^33^. Pax8 is a critical renal lineage transcription factor, thus Cre is diffusely expressed in most renal epithelial cells of the proximal and distal renal tubules, loops of Henle, collecting ducts, and the parietal epithelial cells of Bowman’s capsule following tamoxifen exposure in this model. *SFPQ-TFE3 Pax8 ERT-Cre* (*STP*) mice were born at expected mendelian ratios and survived to adulthood, when they were injected with tamoxifen at 8-12 weeks of age to induce Cre expression. When mice were sacrificed at 3.5 months following tamoxifen, *STP* kidneys were variably enlarged with bilateral solid tumors (**Fig. 2A**), showing strong, diffuse, nuclear TFE3 induction (**Fig. 2B**). Tumor cells were diffusely infiltrative around normal renal tubules, leading to disappearance of the normal cortico-medullary junction, and tumor replacement of a large part of the kidney parenchyma (**Fig. 2B**). Kidney/ body weight ratios and BUN levels were also elevated in some *STP* mice by 3.5 months, though some variability in tumor burden was evident in this inducible model (**Fig. 2C, D**). Histologically, the tumors were comprised of nests of monomorphic epithelioid cells with clear to eosinophilic cytoplasm and large round nuclei (**Fig. 2E**). Though they were invasive, the tumor cells were similar in morphology to intratubular proliferations seen in *STK* kidneys **(Fig. 1I)**, and broadly resembled those seen in human PEComas (**Fig. S3A**). By IHC and immunoblotting, *STP* tumor cells showed strong expression of nuclear TFE3, the canonical TFE3 target, GPNMB, as well as expression of melanocytic and lysosomal markers (PMEL, MelanA and Cathepsin K) commonly upregulated in human MiT/TFE-related neoplasms (**Fig. 2F, G**). In contrast, *STP* tumor cells entirely lacked expression of proximal (LTL) and distal (DBA) renal tubular markers (**Fig. S3B**), as well as cytokeratins, such as type II (CK8) or type I keratins (including CK10, 13, 14, 15, 16, 17 and 18) (**Fig. 2H)**. Notably, *STP* tumor cells showed minimal vimentin expression, and were negative for smooth muscle marker (α-SMA) by IHC (**Fig. 2H)**. Cumulatively, these findings suggested that post-natal expression of *SFPQ-TFE3* within murine renal tubular cells results in highly penetrant, infiltrative epithelioid tumors, with brisk expression of melanocytic/lysosomal markers. Importantly, these markers are characteristic of human MiT/TFE-related tumors, such as the tRCC cases inadvertently included in The Cancer Genome Atlas (TCGA) papillary RCC (KIRP) cohort^34^ **(Fig. 2I)** or tuberous sclerosis (TSC)-related PEComas (angiomyolipomas)^30,35^ (**Fig. 2J**).

**Figure 2:**
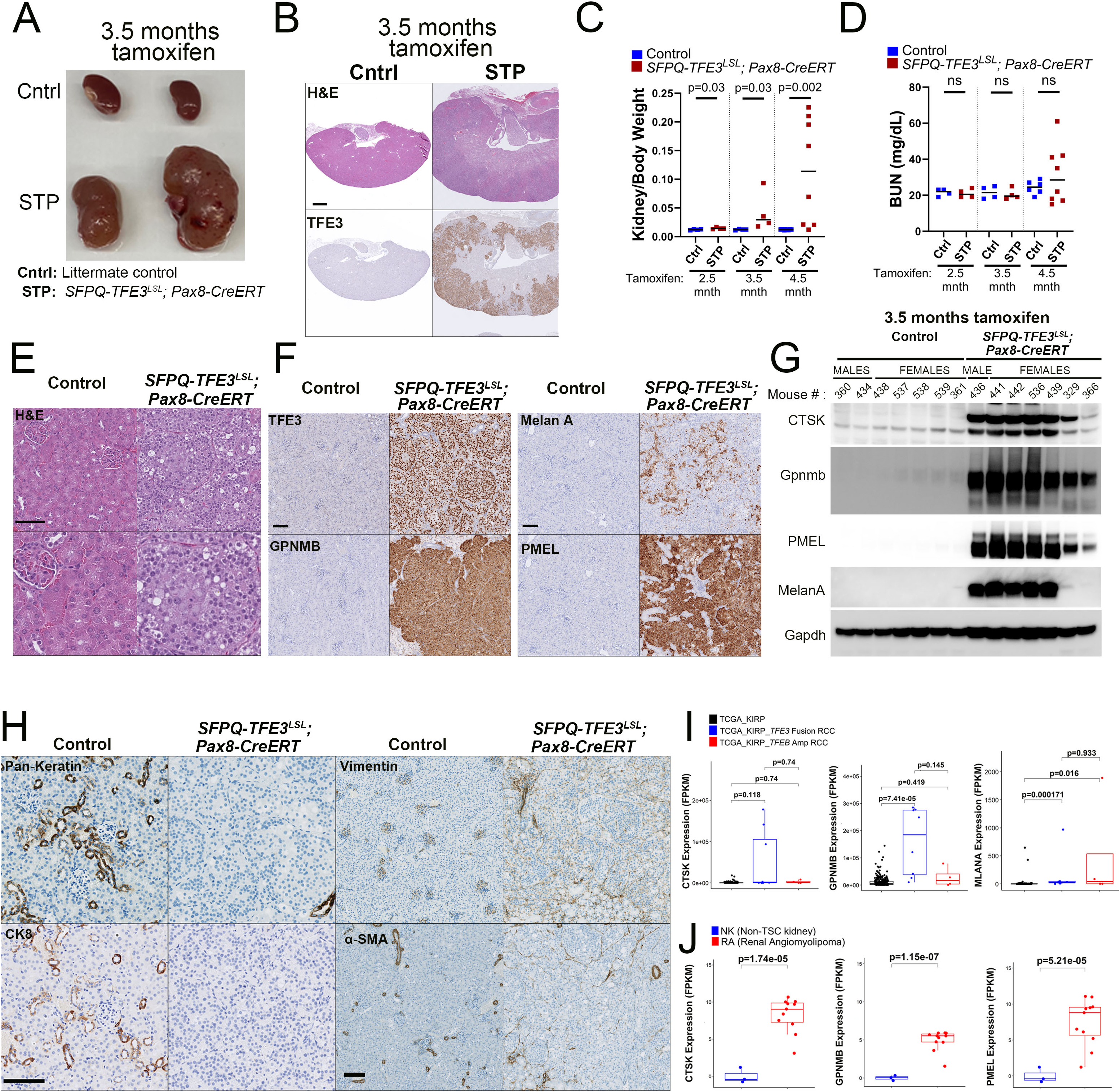
Conditional post-natal, doxycycline-mediated induction of SFPQ-TFE3 in Pax8 Cre-ER^T2^ mice induces renal tumor development. **(A)** Macroscopic images of kidneys from representative *SFPQ-TFE3^LSL^; Pax8-CreERT* transgenic mice and age-matched, littermate controls, sacrificed at 3.5 months following injection of tamoxifen. **(B)** H&E (top row) and TFE3 IHC staining (bottom row) of kidneys from representative *SFPQ-TFE3^LSL^; Pax8-CreERT* transgenic mice and age-matched, littermate controls, sacrificed at 3.5 months following injection of tamoxifen. Scale bar: 1 mm**. (C)** Kidneys to body weight ratios, and **(D)** BUN levels from *SFPQ-TFE3^LSL^; Pax8-CreERT* transgenic mice and age-matched, littermate controls, sacrificed at 2.5, 3.5 or 4.5 months, following injection of tamoxifen. The following numbers were analyzed: a) Kidneys to body weight ratios at 2.5 months [control=4, STP=4], 3.5 months [control=4 and STP=4] and 4.5 months [control=12 and STP=8] and b) BUN levels at 2.5 months [control=4, STP=4)], 3.5 months [control=4 and STP=4] and 4.5 months [control=6 and STP=8]. Data are presented as median. P values by Mann-Whitney test. **(E)** Low (top row) and high (bottom row) magnification images of H&E-stained kidneys from tamoxifen-injected, *SFPQ-TFE3^LSL^; Pax8-CreERT* transgenic mice and age-matched, littermate controls. Scale bar= 100 µm. Representative immunohistochemistry (IHC) for (**F)** TFE3 (top row), GPNMB (bottom row) [left panels], and Melan A (top row), and PMEL (bottom row) [right panels], from tamoxifen-injected, *SFPQ-TFE3^LSL^; Pax8-CreERT* transgenic mice and age-matched, littermate controls. Scale bar= 100 µm. **(G)** Immunoblotting of kidney lysates from tamoxifen-injected, control and *SFPQ-TFE3^LSL^; Pax8-CreERT* transgenic mice at 3.5 months following injection of tamoxifen, for the indicated antibodies. **(H)** Representative immunohistochemistry (IHC) for Pan-Keratin (left panels; top row), CK8 (left panels; bottom row), Vimentin (right panels; top row) and α-SMA (right panels; bottom row) from tamoxifen-injected, *SFPQ-TFE3^LSL^; Pax8-CreERT* transgenic mice and age-matched, littermate controls. Scale bar= 100 µm. **(I)** The gene expression database for papillary RCC (KIRP) from TCGA containing 8 cases of *TFE3* fusion-RCC and 4 cases of *TFEB*-amplified RCC was utilized to compare *Cathepsin K* (*CTSK*), *GPNMB* and *MLANA* gene expression in *TFE3* fusion-RCC and *TFEB*-amplified RCC to the remainder of papillary RCC cases without underlying *TFE3* fusions or *TFEB* amplifications (n=273). Data are represented as a box-and-whisker plot. (*P*-values indicated are by Wilcoxon rank sum test adjusted with multiple comparisons using the false discovery rate (FDR) method). **(J)** Comparison of *Cathepsin K* (*CTSK*), *GPNMB* and *PMEL* expression (FPKM) in RNA seq data from a panel of non-TSC normal kidneys (n=3) and renal angiomyolipomas with *TSC1/2* biallelic loss (n=11)^35^. (*P*-values indicated are by Wilcoxon rank sum test). Source data are provided as a Source data file.

### mTORC1 signaling is activated in murine and human models of SFPQ-TFE3 fusion-driven tRCC

mTOR signaling has previously been shown to be activated in human tRCC samples and pre-clinical models by transcriptomic and proteomic profiling^17,23,24,26,27^, including in a recent mouse model of *ASPSCR1-*TFE3^25^ tRCC, though this signaling pathway was not examined in the previously published *PRCC-TFE3* transgenic mice^3^. We performed gene set enrichment analyses (GSEA)^36^, using Hallmark gene sets^37^ on publicly available gene expression data from the *PRCC-TFE3* mice^3^, and found significant enrichment of genes associated with PI3K/ AKT/ MTOR signaling in the transgenic mice at 7 months compared to controls (**Fig. S4A)**. By immunoblotting, there were increased levels of mTORC1 substrate p-4E-BP1 in *PRCC-TFE3; KSP-Cre* kidney tumor lysates compared to controls, though total 4E-BP1 levels were increased as well (**Fig. S4B)** as has been seen previously in other mouse models with mTORC1 activation in the kidney^19^. We then characterized mTORC1 signaling in the *SFPQ-TFE3* transgenic models. Phosphorylation of canonical mTORC1 substrates and downstream signaling intermediates (p70 S6K, 4E-BP1 and S6) was elevated in P15 *STK* transgenic kidney tumor lysates compared to controls, by immunoblotting (**Fig. 3A**) and IHC (**Fig. 3B**). Levels of phosphorylated canonical mTORC1 substrates and downstream signaling intermediates were also elevated in kidney tumor lysates from 3.5-month *STP* mice compared to treated littermate controls (with substantial increase in total substrate levels seen in these tumors) by immunoblotting (**Fig. 3C**), and similar evidence of mTORC1 activation seen by IHC (**Fig. S4C**). Notably, in the *STP* mice, expression of phosphorylated mTORC1 substrates were specifically increased in the TFE3 (+) tumor cells by IHC, with low expression in surrounding normal kidney (**Fig. 3D**). In addition, levels of nuclear TFEB, an established non-canonical mTORC1 substrate^38^, were also lower in the tubular cells from STK transgenic mice compared to littermate control kidneys by IHC, and a similar pattern was seen in tumor cells from *STP* transgenic mice, compared to internal control surrounding normal renal tubules (**Fig. 3E)**. These findings are consistent with TFEB hyperphosphorylation and cytoplasmic retention due to increased mTOR signaling.

**Figure 3:**
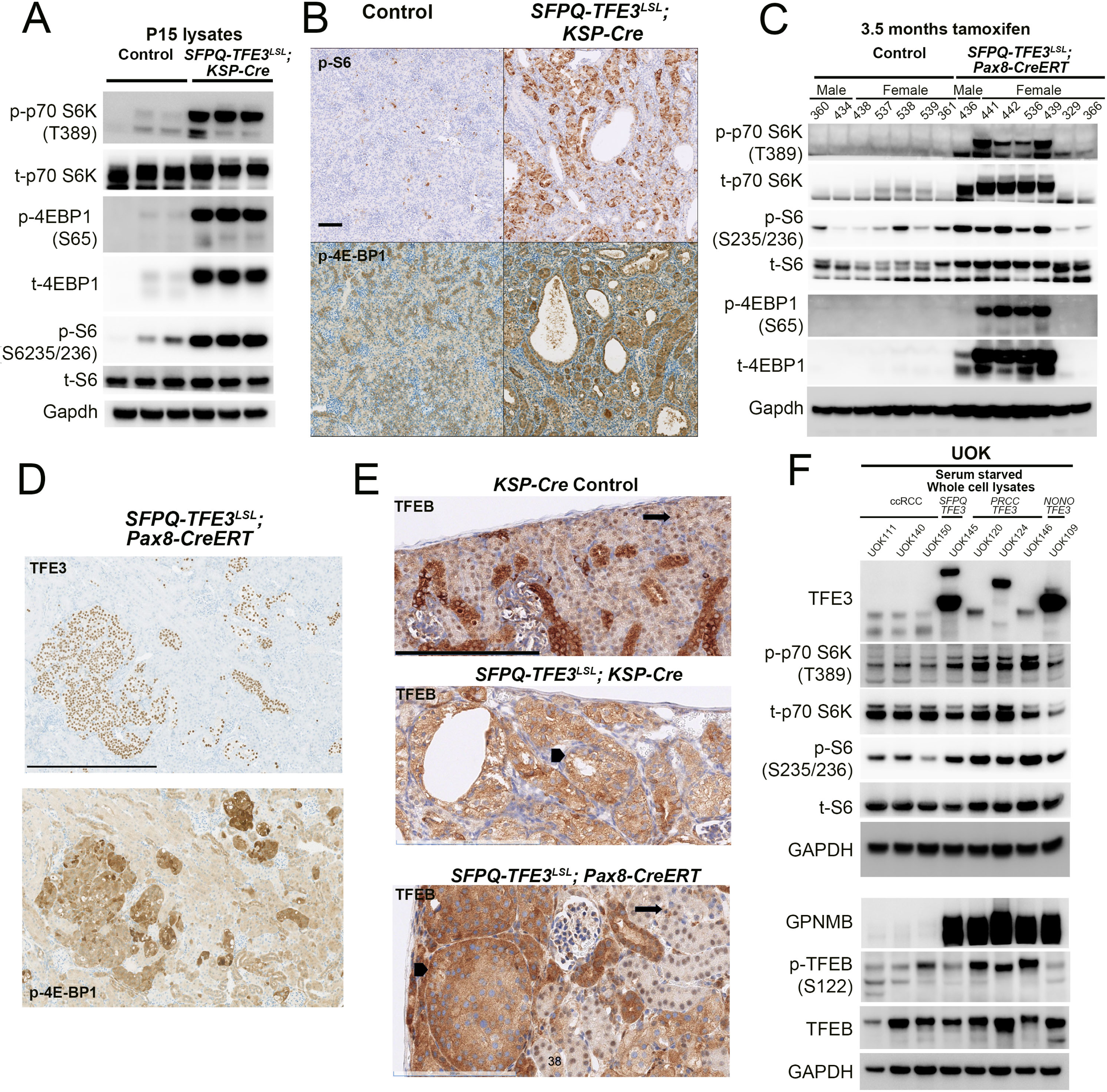
mTORC1 signaling is activated in murine and human models of SFPQ-TFE3 fusion-RCC. **(A)** Immunoblotting of kidney lysates from control and *SFPQ-TFE3^LSL^; Ksp-Cre* transgenic mice at post-natal day 15 for phosphorylation of mTORC1 substrates (p-p70S6K [T389], p-4E-BP1[S65] and p-S6[S235/236]). **(B)** Representative immunohistochemistry (IHC) for p-S6[S235/236] (top row) and p-4E-BP1[T37/46] (bottom row) from age-matched, control and *SFPQ-TFE3^LSL^; Ksp-Cre* transgenic mice at post-natal day 15. Scale bar= 100 µm. **(C)** Immunoblotting of kidney lysates from *SFPQ-TFE3^LSL^; Pax8-CreERT* transgenic mice and age-matched, littermate controls, at 3.5 months following injection of tamoxifen, for phosphorylation of mTORC1 substrates (p-p70S6K [T389], p-4E-BP1[S65] and p-S6[S235/236]). **(D)** Representative immunohistochemistry (IHC) for TFE3 (top row) and p-4E-BP1[T37/46] (bottom row) from tamoxifen-injected, *SFPQ-TFE3^LSL^; Pax8-CreERT* transgenic mice and age-matched, littermate controls, at 2.5 months following injection of tamoxifen. Scale bar= 500 µM. **(E)** Representative immunohistochemistry (IHC) for TFEB from *Ksp-Cre* control mice (top row) and *SFPQ-TFE3^LSL^; Ksp-Cre* transgenic mice (middle row) at post-natal day 15, and from tamoxifen-injected, *SFPQ-TFE3^LSL^; Pax8-CreERT* transgenic mice at 3 months following injection of tamoxifen (bottom row). Arrows indicate nuclear TFEB in normal tubules. Arrowheads indicate cytosolic TFEB in expanded tubules. Scale bar= 200 µM. **(F)** Immunoblotting of lysates from *TFE3* fusion-RCC cell lines [UOK145(SFPQ-TFE3), UOK120,124,146 (PRCC-TFE3) and UOK109(NONO-TFE3)], and ccRCC controls (UOK111, UOK140 and UOK150) for phosphorylation of mTORC1 substrates. Source data are provided as a Source data file.

To validate these findings, we also examined mTORC1 activation in multiple tRCC human cell line models^3^, comparing them to ccRCC cell lines by immunoblotting. Phosphorylation of canonical (p70 S6K) and non-canonical (TFEB) mTORC1 substrates and downstream signaling intermediates (S6) was generally elevated in the tRCC cell lines with *SFPQ-TFE3* (UOK145), *PRCC-TFE3* (UOK120, UOK124, UOK146) and *NONO-TFE3* (UOK109) fusion expression, compared to ccRCC controls (UOK111, UOK140 and UOK150) (**Fig. 3F**). Because comparison of genetically disparate UOK cell lines does not allow examination of the isolated effects of *TFE3* fusions on mTORC1 activation in an isogenic setting, we engineered HEK293 cells with doxycycline-inducible expression of *WT-TFE3*, *S321A-TFE3* (an mTORC1-site phosphodeficient TFE3 point mutant that is constitutively nuclear localized^39^), *SFPQ-TFE3, PRCC-TFE3* and *NONO-TFE3,* using the Flp-In-T-Rex^TM^ system in HEK293 cells. Doxycycline-mediated induction of the variably sized fusion proteins and canonical E-box targets *GPNMB*^3^, *RRAGD*^40^ and *FLCN*^40^ was confirmed by immunoblotting (**Fig. S4D**). Phosphorylation of mTORC1 substrates and downstream signaling intermediates was strongly elevated upon induction of *SFPQ-TFE3* and *PRCC-TFE3* in this system, with subtler induction due to *WT-TFE3*, *S321A-TFE3*, and minimal induction with *NONO-TFE3* (**Fig. S4D)**. To validate these results in a renal tubular cell line, we leveraged a previously described HK2 cell system (human proximal renal tubular epithelial cell line) with doxycycline-inducible expression of *WT-TFE3* and *TFE3* fusion proteins using the rtTA3 (Tet-on) construct^41^. These immunoblotting experiments confirmed increased phosphorylation of mTORC1 substrates and downstream signaling intermediates with expression of *TFE3* fusion proteins in human renal tubular epithelial cells (**Fig. S4E**). A dose-response time course for doxycycline confirmed increased TFEB phosphorylation in HK2 cells with inducible SFPQ-TFE3 expression (**Fig. S4F**). Taken together, these data suggest that MiT/TFE fusion gene expression induces increased mTORC1 activity in human cells.

The mechanism of mTORC1 activation downstream of TFEB expression has previously been ascribed to increased MiT/TFE-mediated transcription of *RRAGD*, an essential component of the Rag GTPase nutrient-sensing complex that regulates amino acid-induced mTORC1 activation at the lysosome^42^ ^18,19,29^. However, to our knowledge, increased *RRAGD* transcription has been documented in only a single cell line model of tRCC^29^. To explore a potential role for *RRAGD,* or its functionally redundant paralog *RRAGC*^43^, in mediating mTORC1 activation in tRCC models, we first examined *RRAGC* and *RRAGD* gene expression in human tRCC specimens^34^ (**Fig. 4A**), where both genes were significantly upregulated in tRCC samples compared to papillary RCC without gene fusion expression. We then examined *RRAGC* and *RRAGD* gene expression in the aforementioned panel of 8 UOK tRCC cell lines, and confirmed increased expression of both genes, most notably *RRAGD*, compared to clear cell RCC control cell lines (**Fig. 4B, 4C**), with similar findings at the protein level by immunoblot (**Fig. 4D**). RagD protein expression was similarly upregulated in HEK293 and HK2 cells upon doxycycline-induced expression of *TFE3* fusion proteins (**Figs. S4D, S4E**). Finally, we examined *Rragc/d* gene expression in *STK* and *STP* transgenic mice kidneys, where it was also increased compared to control kidneys (**Fig. 4E, 4F**).

**Figure 4:**
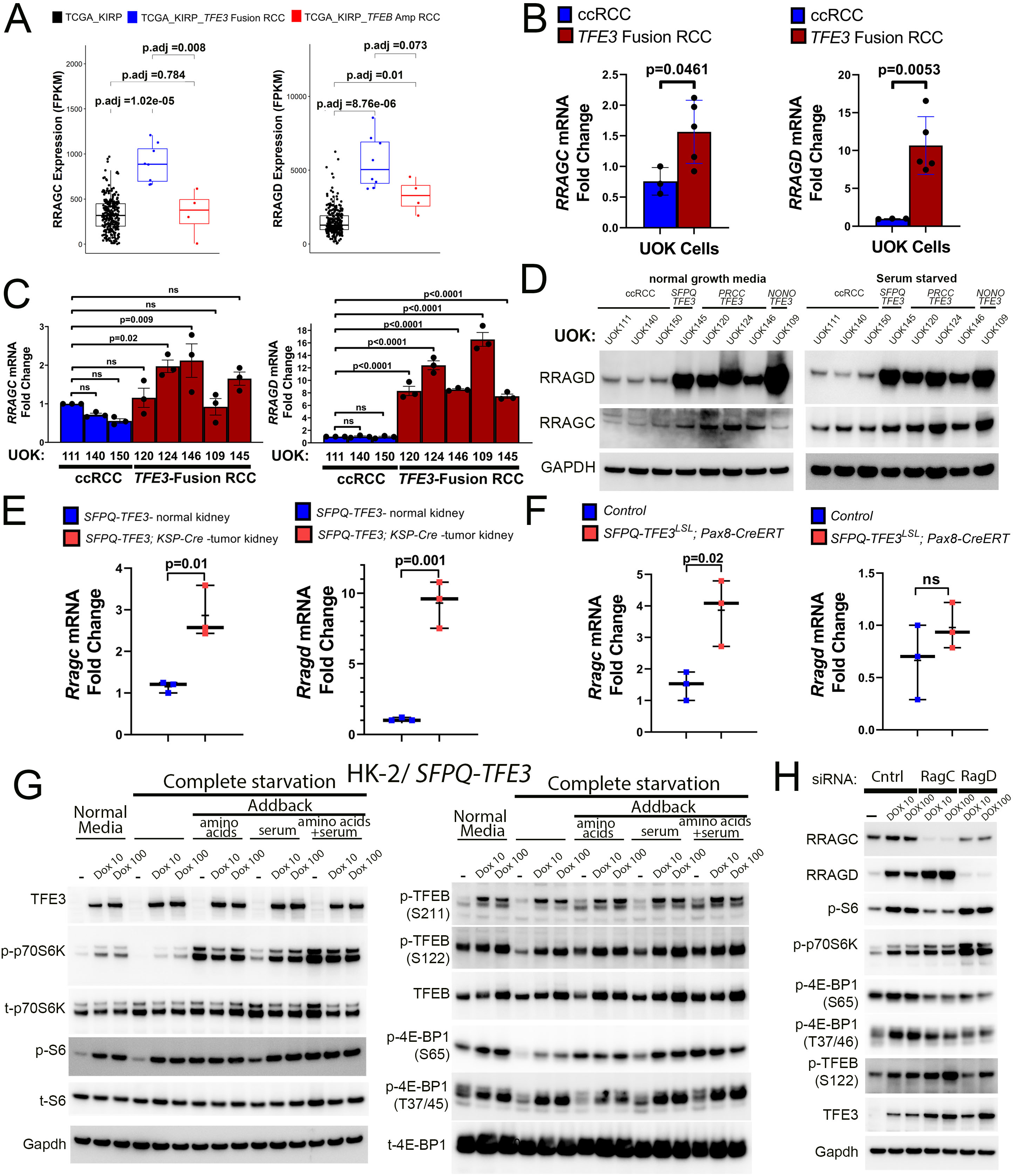
mTORC1 signaling is activated in human models of SFPQ-TFE3 fusion-RCC. **(A)** Comparison of *RRAGC* and *RRAGD* gene expression in *TFE3* fusion-RCC (n=8) and *TFEB*-amplified RCC (n=4) to the remainder of papillary RCC cases without underlying *TFE3* fusions or *TFEB* amplifications (n=273) in the papillary RCC (KIRP) cohort from TCGA. Data are represented as a box-and-whisker plot. (*P*-values indicated are by Wilcoxon rank sum test adjusted with multiple comparisons using the false discovery rate (FDR) method). **(B)** Quantitative real time PCR (qRT-PCR) for *RRAGC* and *RRAGD* transcripts in *TFE3-* fusion RCC cell lines and ccRCC controls (n=3 [ccRCC] and n=5 [*TFE3-* Fusion RCC]; error bars represent SEM; p values by Student’s T-test). **(C)** Quantitative real time PCR (qRT-PCR) for *RRAGC* and *RRAGD* transcripts across individual *TFE3-* fusion RCC cell lines and ccRCC controls (error bars represent SEM; p values by one-way ANOVA). **(D)** Immunoblotting of lysates from *TFE3* fusion-RCC and ccRCC cell lines for RRAGC and RRAGD expression. Quantitative real time PCR (qRT-PCR) for *RRAGC* and *RRAGD* transcripts in kidney lysates from: **(E)** control and *SFPQ-TFE3^LSL^; Ksp-Cre* transgenic mice at post-natal day 15 and **(F)** tamoxifen-injected, *SFPQ-TFE3^LSL^; Pax8-CreERT* transgenic mice at 3.5 months following injection of tamoxifen (n=3, error bars represent SEM; p values by Student’s T-test). **(G)** Immunoblotting of lysates from HK2/*SFPQ-TFE3* cells in normal media, or following nutrient deprivation for 90 min, or following nutrient deprivation followed by stimulation with amino acids, serum or both for 30 min, for the indicated markers. **(H)** Immunoblotting of lysates from HK2/*SFPQ-TFE3* cells following treatment with control, *RRAGC* or *RRAGD* siRNA and the indicated doses of doxycycline for the indicated markers. Source data are provided as a Source data file.

We next examined whether suppression of RRAGC or RRAGD activation via amino acid deprivation or transient *RRAGC* or *RRAGD* silencing was sufficient to suppress mTORC1 signaling in cell line models expressing *TFE3* fusion proteins. In HK-2 cells with inducible *SFPQ-TFE3* expression, amino acid starvation decreased phosphorylation of direct mTORC1 substrates 4E-BP1, p70S6K and TFEB to levels near control cells lacking fusion induction, though relative increased phosphorylation of substrates in cells expressing *SFPQ-TFE3* compared to those without induction remained evident (**Fig. 4G**). Either transient *RRAGC* and/or *RRAGD* knockdown in *SFPQ-TFE3*-expressing cells decreased phosphorylation of 4EB-P1, S6 and/or TFEB to levels seen in control cells, suggesting that they act in a redundant manner, though similar results were not seen for p70S6K phosphorylation (**Fig. 4H**). These data substantiate that RRAGC and RRAGD expression is increased with *TFE3* fusion expression, and transient knockdown of *RRAGC* and/or *RRAGD* is sufficient to suppress phosphorylation of some, but not all, mTORC1 substrates in this context, highlighting the complexities of mTORC1 signaling readouts. Taken together, our results suggest that RRAGC and/or RRAGD may contribute to increased mTORC1 activity in tRCC, but are likely not the only mechanism leading to increased activity of this oncogenic signaling pathway.

### Induction of SFPQ-TFE3 expression in murine renal tubular epithelial cells results in lineage plasticity with silencing of nephric lineage factors Pax8 and Pax2

Although the tumors in our *STP* model were epithelioid and definitively derived from a PAX8-positive cell of origin due to use of the *Pax8 Cre-ER^T2^* model^33^, the lack of keratin and renal tubular marker expression raised the possibility that these tumors more closely resembled malignant PEComas rather than tRCC. To further characterize the lineage of these tumors, we carried out RNA-seq on 15-day *STK* and 3.5-month tamoxifen-treated, *STP* transgenic kidneys, as well as 7-month *PTK* kidneys, and examined differentially expressed genes compared to their respective normal control kidneys (**Supplemental Tables S1-3**). Notably, the core set of differentially expressed genes overlapping between human tRCC and the *ASPSCR1-TFE3* mouse model ^25^ were strikingly enriched in the *STP, STK* and *PTK* transgenic kidney tumors (**Figs. S5A**), highlighting the transcriptional overlap between our mouse models and tRCC. However, gene set enrichment analyses using standard KEGG sets was striking for convergent negative enrichment for the peroxisome and numerous peroxisome-associated metabolic pathways^44^ associated with renal tubular epithelial cells in the *STK*, *STP* and *PTK* models (**Supplemental Tables S4-6**). Consistent with this finding, there was striking negative enrichment of genes associated with renal epithelial cell subsets from the cell type signature gene sets (C8) in *STK* and *STP*, and to a lesser extent in *PTK* transgenic kidneys, consistent with downregulation of keratins and renal tubular markers in these models (**Figs. S5B)**.

To further probe loss of renal tubular cell identity in the *STP* model, we examined expression of core renal lineage transcription factors in tubular cells with or without *SFPQ-TFE3* expression within the same kidney. We used GPNMB expression to identify renal tubules with mosaic *SFPQ-TFE3* fusion transgene expression in mice treated for only two weeks with tamoxifen. Strikingly, nuclear PAX8 expression was conspicuously absent in GPNMB+ tubular epithelial cells, in contrast to adjacent GPNMB-neighboring tubular cells (**Fig. 5A**), indicating that suppression of PAX8 occurs rapidly after *SFPQ-TFE3* fusion expression. Concordant results were seen for keratin (CK8) immunostaining at this timepoint, with CK8 loss occurring specifically in cells with fusion gene expression (**Fig. S6A**). After 3.5 months of tamoxifen treatment, the full-blown tumors in the *STP* model lacked detectable PAX8 (**Fig. 5B**), consistent with loss of keratin in this model **(Fig. 2H**). A more heterogeneous pattern of PAX8 and PAX2 suppression was seen in the *STK* kidneys at P15 (**Fig. 5C**) as well as the *PTK* kidney tumors at 7 months age (**Fig. 5D**) however these differences were highly significant on digital quantification for PAX8 IHC expression (**Fig. 5E, F**), indicating that both *SFPQ-TFE3* and *PRCC-TFE3* expression drives varying degrees of renal lineage factor loss in the murine kidney.

**Figure 5:**
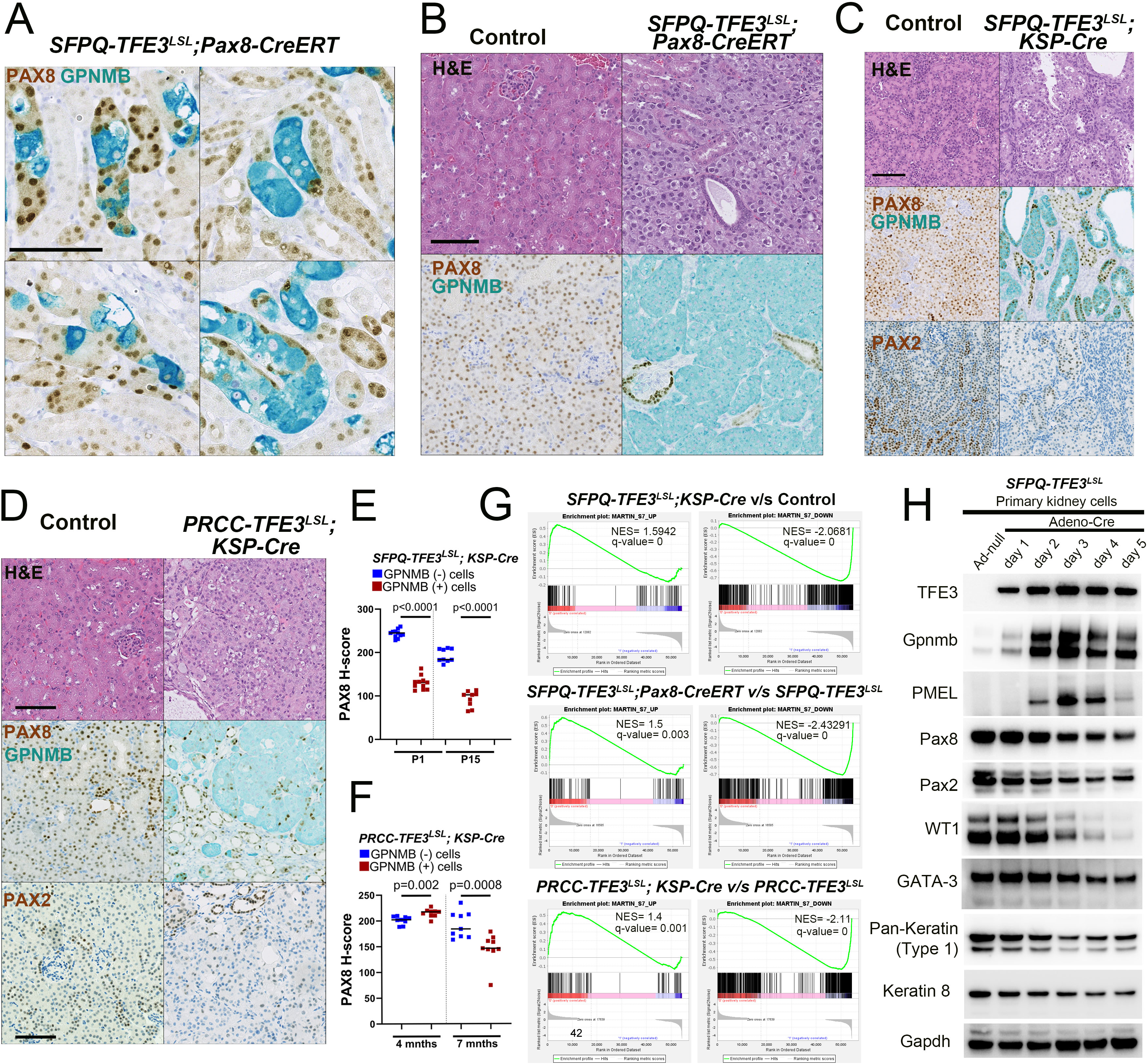
Induction of SFPQ-TFE3 expression in murine renal tubular epithelial cells results in lineage plasticity with silencing of nephric lineage factors, Pax8 and Pax2. **(A)** Dual IHC for PAX8 (brown) and GPNMB (teal) in tamoxifen-injected, *SFPQ-TFE3^LSL^; Pax8-CreERT* transgenic mice and age-matched, littermate controls, at 2 weeks following injection of tamoxifen. Scale bar = 100 µm. **(B)** H&E (top row) and dual PAX8/GPNMB IHC staining (bottom row) of kidneys from representative tamoxifen-injected, *SFPQ-TFE3^LSL^; Pax8-CreERT* transgenic mice and age-matched, littermate controls, at 3.5 months following injection of tamoxifen. Scale bar= 100 µm. **(C)** H&E (top row), dual PAX8/GPNMB IHC staining (middle row) and PAX2 IHC staining (bottom row) of kidneys from representative age-matched, control and *SFPQ-TFE3^LSL^; Ksp-Cre* transgenic mice at post-natal day 15. Scale bar = 100 µm. **(D)** H&E (top row), dual PAX8/GPNMB IHC staining (middle row) and PAX2 IHC staining (bottom row) of kidneys from representative age-matched, control and *PRCC-TFE3^LSL^; Ksp-Cre* transgenic mice at 7 to 9 months. **(E)** Digital quantification of mean nuclear PAX8 H-scores in GPNMB-cells (blue) and GPNMB+ cells (red) at day 1 and day 15 (Fix Figure), from experiments in **C**, as depicted by scatter plots, with the central line denoting the median. The number of biological replicates analyzed for both, GPNMB – and + cells were n=11 (day 1) and n=9 (day 15). Statistical analyses were performed using two-tailed Mann-Whitney test. **(F)** Digital quantification of mean nuclear PAX8 H-scores in GPNMB-cells (blue) and GPNMB+ cells (red) at 4 months and 7 months, from experiments in **D**, as depicted by scatter plots, with the central line denoting the median. The number of biological replicates analyzed for both, GPNMB – and + cells were n=9 at 4 and 7 months. Statistical analyses were performed using two-tailed Mann-Whitney test. **(G)** Gene Set Enrichment Analysis (GSEA) comparing 15-day *STK* (top panels), 3.5-month tamoxifen-treated, *STP* (middle panels) and 7-month *PTK* (bottom panels), transgenic kidneys and their controls, for genes differentially expressed in renal TSC-related PEComas from^35^. **(H)** Immunoblotting of lysates from primary renal tubular epithelial cells from *SFPQ-TFE3^LSL^*transgenic mice treated with control or Cre-recombinase expressing adenovirus *in vitro* for the indicated markers. Source data are provided as a Source data file.

The *PAX8* transcription factor hub drives a core regulon of target genes in the kidney to promote proximal tubular epithelial cell fate, including *GATA3*, *LHX1* and *WT1*, among others^45^, and we confirmed these targets in HK2 proximal tubular epithelial cells following *PAX8* knockdown using immunoblotting and qRT-PCR (**Fig. S6B, S6C**). Loss of PAX8 phenocopied expression patterns in renal TSC-related PEComas (angiomyolipomas), which show striking downregulation of *PAX8* or *PAX2* and their core target genes (*GATA3, WT1, LHX1,*)^35,45^ compared to surrounding kidney parenchyma. (**Fig. S6D**). Indeed, expression of GATA3 and pan (type I) keratin was also markedly decreased in *STP* tumors by 3.5 months after tamoxifen treatment, consistent with loss of PAX8 (**Fig. S6E**) and *STK, STP* and *PTK* transgenic kidneys showed significant enrichment of both up- and downregulated genes from renal TSC-related PEComas compared to controls (**Fig. 5G**)^35^. Finally, to explore the temporal dynamics of downregulation of *PAX8*, *PAX2* and downstream target genes following *SFPQ-TFE3* fusion expression, we leveraged cultured primary renal tubular epithelial cells from *ST* mice, treated with empty or Cre-recombinase-expressing adenovirus *in vitro.* By immunoblotting, expression of PAX8, PAX2 and their transcriptional targets GATA3 and WT1, as well as pan-keratin (Type 1) were decreased within two days of *SFPQ-TFE3* induction (**Fig. 5H)**. Taken together, both *SFPQ-TFE3* and *PRCC-TFE3* fusion expression are accompanied by downregulation of renal lineage transcription factors and their downstream targets *in vivo* and *in vitro* in the murine kidney. This finding is most dramatic in the *STP* model, where resulting tumors are best classified as renal epithelioid malignant PEComas, based on total loss of PAX8, PAX2, cytokeratin and renal tubular marker expression, with accompanying upregulation of melanocytic and lysosomal markers. Since *SFPQ-TFE3* expression is limited to PAX8-expressing cells in the *STP* model due to use of the *Pax8 Cre-ER^T2^* model, this model serves as a lineage tracing experiment, thereby substantiating renal tubular epithelial cells as the cell of origin for a TFE3 fusion-driven murine PEComa model.

### Induction of SFPQ-TFE3 expression in human renal tubular epithelial cells results in lineage plasticity with silencing of renal epithelial lineage factors PAX8 and PAX2

To validate our murine results in human systems and to further test whether findings were conserved across different *TFE3* fusion partners, we next examined expression of PAX8 and PAX2 in the HK2 proximal renal epithelial cell line with doxycycline-inducible expression of common *TFE3* fusions. Similar to findings in our mouse models, PAX8 and PAX2 protein expression were most strikingly and specifically downregulated with induction of *SFPQ-TFE3* and this was accompanied by robust upregulation of the melanocytic marker PMEL (**Fig. 6A**). Notably, WT-TFE3 over-expression did not affect PAX8 or PAX2 expression, while *PRCC-TFE3* induction was accompanied by a mild suppression of PAX2, without discernable effects on PAX8 expression. PAX8 nuclear localization was also significantly decreased in HK2/*SFPQ-TFE3* cells by nuclear-fraction immunoblotting (**Fig. 6B**) and immunofluorescence (**Fig. 6C**). These findings were corroborated at the gene expression level, where *PMEL* and *GPNMB* expression were dramatically increased upon *SFPQ-TFE3* induction (**Fig. 6D**), and accompanied by decreased mRNA expression of *PAX8*, *PAX2* and the downstream transcriptional targets of these renal lineage transcription factors, including *LHX1, HNF1B* and *GATA3* (**Fig. 6E**). The downregulation of PAX8 with doxycycline induction of *SFPQ-TFE3* was also rescued in HK2/*SFPQ-TFE3* cells stably expressing TFE3-targeting CRISPR sgRNAs (**Fig. 6F**), providing evidence that PAX8 silencing is due to *SFPQ-TFE3* expression and not a confounding effect of doxycycline treatment in this system.

**Figure 6:**
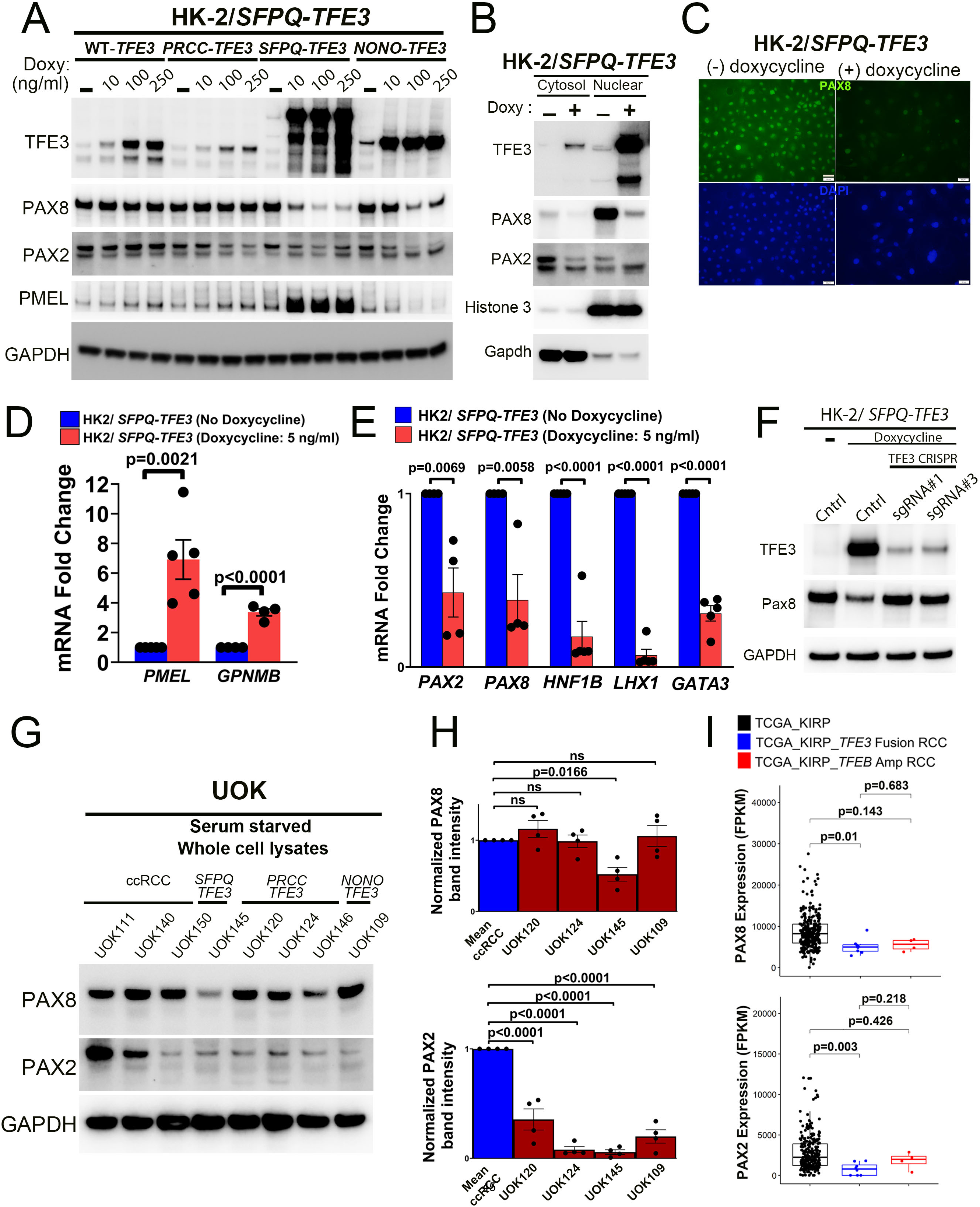
Induction of SFPQ-TFE3 expression in human renal tubular epithelial cells results in lineage plasticity with silencing of nephric lineage factors, PAX8 and PAX2. **(A)** Immunoblotting of HK2 proximal tubular epithelial cells with doxycycline-inducible expression of *WT-TFE3* and *TFE3* fusion-proteins, for the indicated markers. Cells were treated with the indicated doses of doxycycline for 72 hrs, prior to lysis and immunoblotting. **(B)** Immunoblotting of cytosolic and nuclear fractions of HK2 cells with doxycycline-inducible expression of *SFPQ-TFE3* for the indicated markers. **(C)** Indirect immunofluorescence for PAX8 in HK2 cells with doxycycline-inducible expression of SFPQ-TFE3. (D) Quantitative real time PCR (qRT-PCR) for *PMEL* and *GPNMB* in HK2 cells with doxycycline-inducible expression of *SFPQ-TFE3* (n>3, error bars represent SEM; p values by Student’s T-test). **(E)** Quantitative real time PCR (qRT-PCR) for *PAX2, PAX8, HNF1B, LHX1* and *GATA3* in HK2 cells with doxycycline-inducible expression of *SFPQ-TFE3* (n>3, error bars represent SEM; p values by Student’s T-test). **(F)** Immunoblotting of HK2 cells with doxycycline-inducible expression of *SFPQ-TFE3* with CRISPR-Cas9-mediated genomic inactivation of *TFE3,* for TFE3 and PAX8. Two clones representing two unique guide RNAs targeting *TFE3* are shown. **(G)** Immunoblotting of lysates from *TFE3*-fusion RCC cell lines [UOK145(SFPQ-TFE3), UOK120,124,146 (PRCC-TFE3) and UOK109(NONO-TFE3)], and ccRCC controls (UOK111, UOK140 and UOK150) for PAX8 and PAX2. **(H)** Normalized densitometric quantifications for PAX8 and PAX2 immunoblots, from experiments in G (n=4, error bars represent SEM; p values by one-way ANOVA). **(I)** Comparison of *PAX8* and *PAX2* gene expression in *TFE3* fusion-RCC (n=8) and *TFEB*-amplified RCC (n=4) to the remainder of papillary RCC cases without underlying *TFE3* fusions or *TFEB* amplifications (n=273) in the papillary RCC (KIRP) cohort from TCGA. Data are represented as a box-and-whisker plot. (*P*-values indicated are by Wilcoxon rank sum test adjusted with multiple comparisons using the false discovery rate (FDR) method). Source data are provided as a Source data file.

Though tRCC is distinguished from PEComa by its retention of detectable keratin and PAX8 expression in clinical practice, the downregulated expression of PAX8 expression in the *PTK* tRCC model and the frequent underexpression of cytokeratin and EMA expression seen in human tRCC ^17^ led us to test whether there is partial *PAX8* and *PAX2* loss in human tRCC samples that had not been previously appreciated. We first examined the series of patient-derived tRCC cell lines, where UOK145 cells expressing *SFPQ-TFE3* showed a striking reduction in PAX8 protein expression compared to ccRCC cell lines, but nearly all tRCC cell lines showed some decrease in PAX2 expression compared to the average expression seen in ccRCC lines (**Fig. 6G, H**). Leveraging the TCGA papillary RCC (KIRP) cohort^34^, mRNA expression levels of *PAX2* and *PAX8* were significantly decreased in *TFE3* fusion-RCC cases, compared to ccRCC (**Fig. 6I),** while expression levels of related family members *PAX5* and *PAX6* remained unaltered (**Fig. S6F**), though sample numbers were too small to examine the effects of specific fusions on *PAX8* and *PAX2* expression. Cumulatively, our results in human samples indicate that *SFPQ-TFE3* expression is associated with particularly potent suppression of renal lineage transcription factor expression compared to other common *TFE3* fusion genes, with resulting lineage plasticity towards a PEComa phenotype, while other fusion genes have a milder effect. These findings are entirely consistent with the fact that *SFPQ* is the most common *TFE3* fusion partner in human PEComas and further support the fidelity of current MiT/TFE mouse tumor models, where the *PRCC-TFE3* model shows partial retention of PAX8 and epithelial features consistent with tRCC^3^, while our *SFPQ-TFE3* model most closely approximates a malignant epithelioid PEComa phenotype.

### mTOR inhibition rescues PAX2 and PAX8 expression and activation in human vitro models of SFPQ-TFE3 expression

Finally, we examined whether mTOR signaling activity reinforces renal tubular cell lineage plasticity downstream of *SFPQ-TFE3* fusion expression. Leveraging our *in vitro* models for pharmacologic and genetic mTOR modulation, we first asked whether suppression of mTOR signaling concurrent with induction of fusion TFE3 expression in HK2/*SFPQ-TFE3* cells affected PAX2/PAX8 expression or MiT/TFE transcriptional target gene expression. Treatment of HK2/*SFPQ-TFE3* cells with the mTOR kinase inhibitor Torin1, rescued PAX2 and PAX8 protein expression in a dose-dependent fashion and simultaneously downregulated expression of the PEComa marker PMEL, as well as multiple MiT/TFE-regulated proteins (LC3A/B, RRAGD, RAB7A, and GPNMB), by immunoblotting (**Fig. 7A, B**, **S7A**, **B**). These effects were reproduced by pre-treatment of HK2/*SFPQ-TFE3* cells with *RHEB* siRNA to knockdown a key component of the mTOR complex prior to doxycycline induction (**Fig. 7C, D, S7B**), providing genetic confirmation. *SFPQ-TFE3*-induced downregulation of keratin expression was also completely rescued by Torin1 and *RHEB* siRNA (**Fig. S7B**). Strikingly, mTOR inhibition concurrent with doxycycline induction via either Torin1 or *RHEB* siRNA was associated with significantly lower *SFPQ-TFE3* fusion protein expression, and this was particularly evident for *RHEB* siRNA (**Fig. 7A-D, S7A, B**) and mildly dose-dependent for Torin1 treatment (**Fig. S7A**). Accordingly, mTORC1 inhibition with Torin1 rescued PAX8 nuclear localization in doxycycline-treated HK2/*SFPQ-TFE3* cells, by nuclear-fraction immunoblotting (**Fig. 7E**) and immunofluorescence (**Fig. 7F**), with both experiments demonstrating a concurrent decrease in nuclear *SFPQ-TFE3* levels. The dose-dependent decrease in *TFE3* fusion protein expression upon Torin1 and *RHEB* siRNA treatment was generalizable to other cell line systems and *TFE3* fusion partners examined by immunoblotting, including HEK293 cells with doxycycline-inducible expression of *SFPQ-TFE3* (**Fig. S7C**) or *PRCC-TFE3* (**Fig. S7D**) as well as UOK120 and UOK124 cells (**Fig. S7E**) constitutively expressing *PRCC-TFE3*. Finally, Torin1 treatment of HK2/*SFPQ-TFE3* cells downregulated gene expression of MiT/TFE transcriptional targets (GPNMB and PMEL) (**Fig. 7G**), rescued *PAX2* and *PAX8* gene expression and also upregulated expression of PAX2/8-regulated transcripts (*LHX1, GATA3, HNF1B*), by qRT-PCR (**Fig. 7H**), indicating that PAX2/8 regulation by *SFPQ-TFE3* is mediated at the gene expression level, downstream of elevated mTOR activity. Taken together, our findings indicate that mTOR signaling activation occurs downstream of *SFPQ-TFE3* gene fusion expression, and this signaling reciprocally positively regulates *SFPQ-TFE3* fusion levels and transcriptional activity. This positive feedback loop reinforces a lineage switch in renal tubular cells towards a PEComa cell phenotype, characterized by downregulated transcription of renal lineage transcription factors, in a manner that is reversible by genetic or pharmacologic mTOR inhibition (**Fig. 8**).

**Figure 7:**
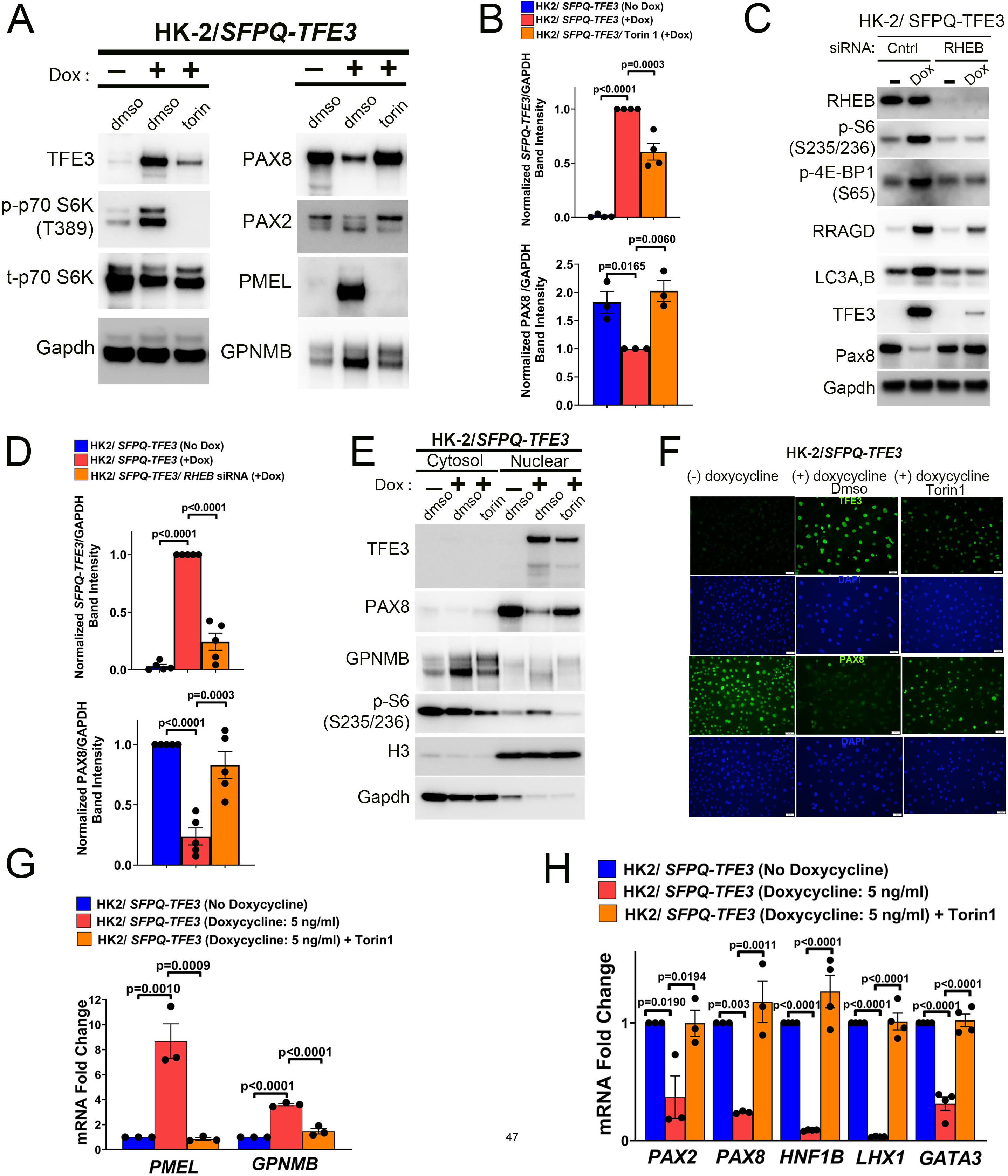
mTORC1 inhibition rescues PAX2 and PAX8 expression and activation in human vitro models of SFPQ-TFE3 expression. **(A)** Immunoblotting of HK2 cells with doxycycline-inducible expression of *SFPQ-TFE3,* following treatment with doses of the mTOR kinase inhibitor torin1, for the indicated antibodies. **(B)** Densitometry quantification of normalized SFPQ-TFE3 and PAX8 expression, in HK2 cells with doxycycline-inducible expression of *SFPQ-TFE3* following treatment with torin1, from experiments in (A) (n>3, error bars represent SEM; p values by one-way ANOVA with Dunnett’s test for multiple comparisons). **(C)** Immunoblotting of HK2 cells with doxycycline-inducible expression of *SFPQ-TFE3,* following treatment with *Rheb* siRNA, for the indicated antibodies. **(D)** Densitometry quantification of normalized SFPQ-TFE3 and PAX8 expression, in HK2 cells with doxycycline-inducible expression of *SFPQ-TFE3* following treatment with *Rheb* siRNA from experiments in (C) (n>3, error bars represent SEM; p values by one-way ANOVA with Dunnett’s test for multiple comparisons). **(E)** Immunoblotting of cytosolic and nuclear fractions of HK2 cells with doxycycline-inducible expression of *SFPQ-TFE3,* following treatment with the mTOR kinase inhibitor torin1 for the indicated markers. **(F)** Indirect immunofluorescence for TFE3 and PAX8 in HK2 cells with doxycycline-inducible expression of *SFPQ-TFE3,* following treatment with the mTOR kinase inhibitor torin1. **(G)** Quantitative real time PCR (qRT-PCR) for *PMEL* and *GPNMB* in HK2 cells with doxycycline-inducible expression of *SFPQ-TFE3,* following treatment with the mTOR kinase inhibitor torin1 (n=3, error bars represent SEM; p values by one-way ANOVA with Dunnett’s test for multiple comparisons). **(H)** Quantitative real time PCR (qRT-PCR) for *PAX2, PAX8, HNF1B, LHX1* and *GATA3* in HK2 cells with doxycycline-inducible expression of *SFPQ-TFE3,* following treatment with the mTOR kinase inhibitor torin1 (n=3, error bars represent SEM; p values by one-way ANOVA with Dunnett’s test for multiple comparisons). Source data are provided as a Source data file.

**Figure 8.**
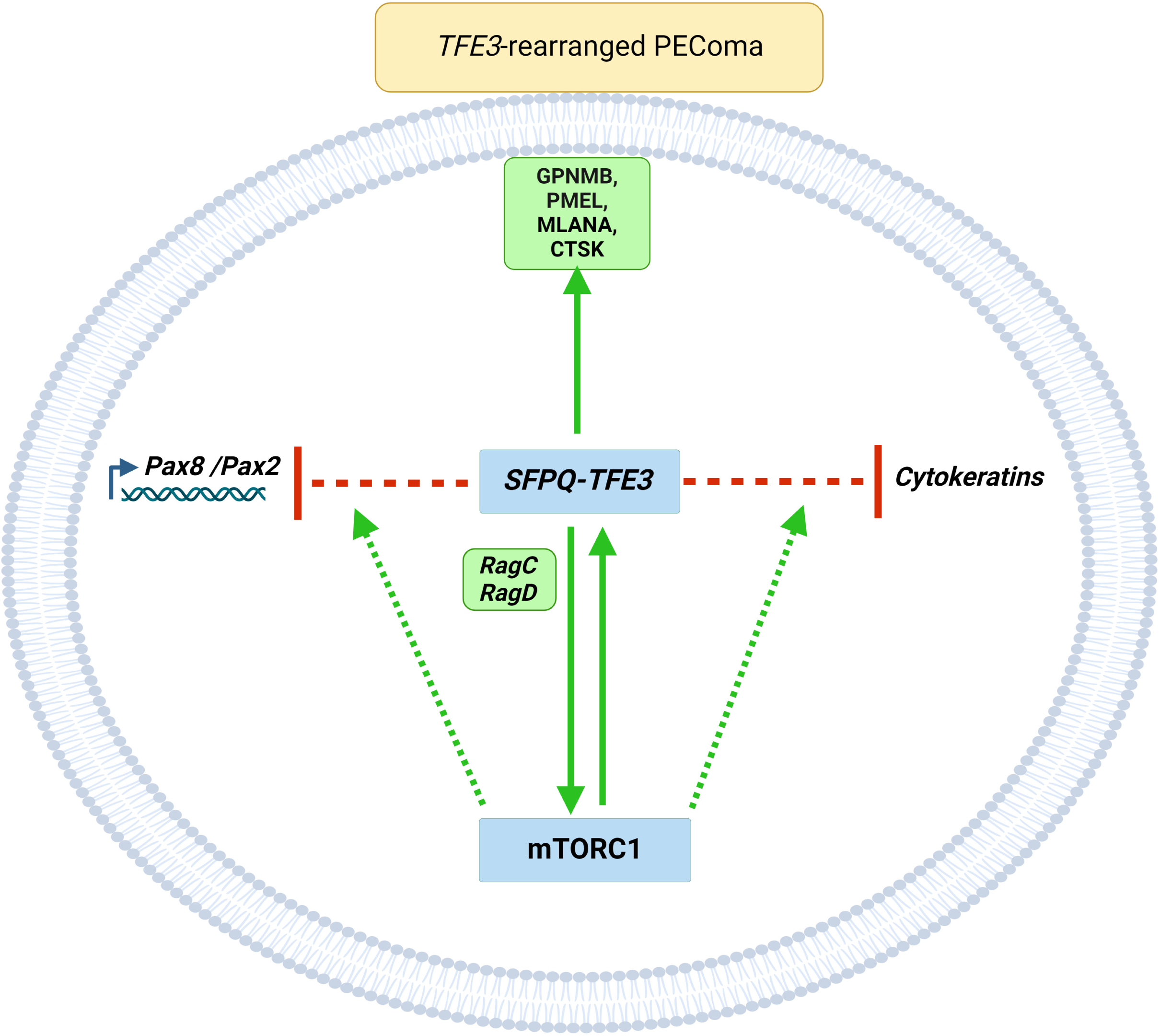

## Discussion

tRCC is characterized by minimal somatic alterations^46^ other than MiT/TFE-fusion events, which are considered necessary and sufficient to induce renal tumor development. However, there is considerable phenotypic heterogeneity in renal tumors driven by different *TFE3*-fusion proteins, in part due to varying functions of the fusion partners, underscoring a need to develop fusion-specific models to study the disease. Here, we provide the first ever characterization of constitutive and inducible transgenic mouse models of the *SFPQ-TFE3* fusion oncoprotein-mediated tumorigenesis in the kidney. Similar to previous studies with constitutive expression of *ASPSCR1-TFE3*^25^, wild-type TFEB^47^, or inactivation of *FLCN* ^48,49^ during renal development, expression of SFPQ-TFE3 using Ksp-Cadherin-Cre (*SFPQ-TFE3^LSL^; Ksp-Cre* mice; *STK*) resulted in disrupted renal development and renal insufficiency, culminating in early neonatal death. In contrast, constitutive induction of *PRCC-TFE3* resulted in a cystic renal phenotype that was compatible with life and resulted in delayed tumorigenesis^3^. These variations across constitutive models likely reflect a combination of factors: **a)** the timing of gene perturbation, since the pathological consequences of gene inactivation are strongly influenced by the developmental status of the kidney, as has previously been well described in the case of *PKD1* deletion^50^, **b)** the tubular origin of the Cre driver used to express the transgene, and perhaps most significantly, **c)** the fusion gene subtype.

In contrast to constitutive, early expression using Ksp-Cadherin-Cre, tamoxifen-induced, postnatal induction of SFPQ-TFE3 [*SFPQ-TFE3* Pax8 Cre-ER^T2^ mice, *STP*], resulted in large, highly penetrant, infiltrative, epithelioid tumors without renal developmental defects. Strikingly, and previously unreported in other fusion-*TFE3* models, *SFPQ-TFE3* expression in both constitutive and inducible models was sufficient to downregulate expression of renal lineage transcription factors PAX8 and PAX2. While, the *ASPSCR1-TFE3* fusion model resulted in extrarenal PEComas^25^, PAX8 was widely expressed in the renal tumors in this model, suggesting that these are best classified as tRCC, though quantification of PAX8 levels relative to normal kidney was not performed. Similarly, the *PRCC-TFE3* model showed retained (though, as we demonstrate, focally low) expression levels of PAX8, consistent with their classification as tRCC^3^. In contrast, postnatal induction of *SFPQ-TFE3* in all PAX8-expressing tubular cells resulted in non-epithelial renal tumors that reproduced the morphology and immunophenotype of human PEComas without PAX8 expression, underscoring the more pronounced role of *SFPQ-TFE3* in driving lineage plasticity, and consistent with the high prevalence of this fusion in human PEComas^30^. Taken together, these models suggest that TFE fusions are associated with a continuum of renal lineage factor downregulation, with *SFPQ-TFE3* at the furthest extreme, and future work will examine what mechanisms might underlie these differences.

The origin and developmental mechanisms that give rise to human PEComas have remained elusive thus far. Cell line models of *TSC* loss^51^ or *TFE3*-fusions^52^ are frequently characterized by cellular senescence, and this has hampered the identification of a potential cell of origin. *SFPQ-TFE3* expression in our inducible *in vitro* models also rapidly induced senescence and apoptosis, with upregulation of p21 and p16 (data not shown). Renal PEComas are most commonly driven by mTORC1 hyperactivation resulting from biallelic inactivation of *TSC1* or *TSC2*, with both constitutional^14,53^ as well as somatic mutagenesis in the *TSC1/2* complex^15,54^. Mechanistically, we and others have shown that mTORC1 activation leads to constitutive activation of *TFEB* and *TFE3*, explaining why TSC1/2 loss and TFE3 fusion expression are mutually exclusive drivers of human PEComas^18,19^. However, homozygous and/or heterozygous deletion of *TSC1* or *TSC2* generally does not induce the formation of renal AML in mice^55,56^, although one previous mouse model of mosaic *TSC1* deletion developed renal mesenchymal lesions that resembled human PEComas, with upregulated melanocytic marker expression^57^. Mechanistically, there is evidence that mTORC1 can itself modulate stemness and plasticity^58^: PEComas in TSC have shown resemblance to the embryonic kidney^35^ and mTORC1 activation via *TSC1/2* loss in nephron progenitors drives pluripotency along multiple (renal epithelial, stromal, and glial) lineages in the developing kidney^15,53^. mTORC1 activation also induces an EMT program in developing renal organoids which is known to overlap with the stemness transcriptional program, thus offering some insight into their pathogenesis^53,59^. Similar to tumors with *TSC1/2* loss, constitutively nuclear-localized, wild-type TFE3 can also function as a cell-fate switch and prevents lineage commitment and differentiation of embryonic stem cells (ESCs), thereby maintaining a state of pluripotency^60^. Our findings suggest that TFE3 fusion proteins may be the most potent fusion at initiating and maintaining plasticity in the developing and postnatal renal tubular epithelium, with more dramatic and complete trans-differentiation compared to that induced by wild-type TFE3 or *TSC1/2* loss. Future ChIP-seq and ATAC-seq experiments will help to explore how the cistrome landscape and chromatin architecture are uniquely impacted by wild-type TFE3 compared to individual *TFE3*-fusions.

The paired-box transcriptional regulators *PAX2* and *PAX8* are closely related members of the PAX family of genes that are co-expressed in the developing kidney, and are vital for specifying the nephric lineage, survival and morphogenesis during kidney development^61,62^. PAX2 and PAX8 share DNA-recognition motifs and are functionally redundant in several aspects of kidney development such as branching morphogenesis, mesenchymal to epithelial transition and nephron differentiation^63–65^. PAX proteins, via interactions with specific co-factors, are known to act as epigenetic regulators to imprint active or repressive histone marks on chromatin, thus driving renal epithelial cell fate^62^. Accordingly, *Pax2* and *Pax8* double mutant mice exhibit a complete lack of kidney formation, due to an increase in apoptosis during development^63,64^. This developmental reliance on the PAX proteins may in part explain the renal developmental defects in our STK model, where these proteins were dramatically downregulated.

In contrast, the role of the PAX proteins in adult kidneys and renal tumorigenesis is not fully understood. In the adult human kidney, PAX8 is expressed in renal epithelial cells in all nephron segments, including the proximal tubules, renal papillae and in the parietal cells of Bowman’s capsule ^66^, while PAX2 is mainly expressed in the collecting ducts, medulla and renal papillae, where they function to promote water/solute homeostasis and osmotic tolerance^67^ or resistance to acute ischemic kidney injury^68,69^ in a redundant manner. Increased tumor-associated expression of PAX2 and/or PAX8 is observed in an overwhelming majority of primary and metastatic clear, papillary and chromophobe renal tumors ^66,70^ and correlates with proliferation index and metastatic disease^71^. Ectopic PAX2 expression is associated with PKD, RCC and Wilms’ tumor^72^ and PAX8 is a presumed oncogene in ccRCC^10,11^, suggesting that many renal tumors exhibit a lineage-dependence on PAX2/8 expression to drive tumorigenesis. tRCC retains expression of PAX8 by definition, however high expression of melanocytic markers and low expression of keratins and EMA corroborates our finding that these tumors downregulate PAX8 relative to the normal kidney ^17^. Similarly, emerging evidence from ccRCC suggests, that a subset of the PAX8 regulon comprised of core target genes (*WT1, LHX1, GATA3*) is selectively downregulated with disrupted terminal epithelial differentiation^45^. Furthermore, HNF1B - which is a direct transcriptional target of PAX8^73^ - is often silenced in PCKD^74^ or chRCC^75^, indicating that loss of PAX8 expression or transcriptional activity is compatible with renal tumorigenesis, and may indeed drive lineage plasticity. Intriguingly, PAX2 and PAX8, in addition to specifying nephron identity, also prevent trans-differentiation along alternate lineages: PAX2 loss initiates a cell-fate switch into renal interstitial cells^76^, and PAX8 loss impedes the MET process and promotes a mesenchymal phenotype in kidney progenitors^77^, consistent with our findings that *SFPQ-TFE3*-induced downregulation of PAX2 and PAX8 in kidney cells may act as a trigger to initiate lineage plasticity and loss of epithelial markers.

While studies on the pathogenesis of PEComas have largely established the role of the 2 main genomic drivers (*TSC1/2* loss and *TFE3* fusions), recent work has further characterized their interdependence. *TSC1/2* loss via mTORC1 activation drives *TFEB/TFE3* activity in cellular^78^ and mouse models^19^ of TSC, via *FLCN* inactivation^18^. Correspondingly, mTORC1 activation, which was hitherto known to be regulated by multiple well-characterized inputs, largely restricted to growth factor signaling (TSC/Rheb complex)^79^, nutrient sensing (RAG GTPases)^42^ and intra-lysosomal sensing (v-ATPase/ Ragulator complex)^80,81^, is now understood to be also regulated by the MiT/TFE factors, via transcriptional induction of the CLEAR gene target *RRAGD* that drives lysosomal recruitment and activation of mTORC1^18,19,29^. While induction of RagD was more robust compared to RagC in our *in vitro* models, in HK2/*SFPQ-TFE3* cells, transient silencing of either *RRAGC* or *RRAGD* was sufficient to decrease 4E-BP1 phosphorylation, while p-TFEB was only decreased by *RRAGD* silencing. These findings are consistent with a recent study that highlighted a RagC/D mTORC1 dimer code determining substrate specificity^82^. Interestingly, activating mutations in *RRAGD* have been associated with constitutive mTORC1 signaling in kidney tubulopathies^83,84^, suggesting that RRAGD overexpression alone may be insufficient to drive mTORC1. Furthermore, in tRCC, in addition to *RRAGD*, multiple regulators of lysosomal mTORC1 signaling (*RHEB*^46^, *RRAGB*^25^, *RRAGC*^25,46^ *FNIP1/2*^25^), are also transcriptionally activated, suggesting a more complex mechanism of mTORC1 activation may be at work. In support of this, gene expression of *RHEB* and multiple *V-ATPASE* components were significantly upregulated in tRCC samples compared to papillary RCC without gene fusion expression in The Cancer Genome Atlas (TCGA) papillary RCC (KIRP) cohort^34^ (data not shown). Further studies are required to determine the role of the numerous upstream signaling inputs driving mTORC1 activation in tRCC.

Underscoring the importance of mTORC1 as an oncogenic driver, multiple groups have reported its activation in pre-clinical models of tRCC. However, studies thus far have demonstrated variable anti-tumor effects of mTOR inhibitors in cell line/xenograft models^26,27,85^, transgenic mouse models^25^ and in patients for tRCC^86–88^ and PEComas^89^, with mTOR kinase inhibitors conferring a greater benefit^85^ than rapalogs^25^ in pre-clinical studies. These findings may potentially be due to robust downregulation of fusion-*TFE3* expression with mTOR kinase inhibitors, as we noted in multiple cell lines expressing *TFE3* fusions. However, unlike wild-type MiT/TFE factors that are inhibited by mTORC1-mediated phosphorylation^90^, *TFE3*-fusion proteins have until now, been considered constitutively nuclear localized and therefore unrestrained by mTORC1^1^. Interestingly, Torin 1 induced accelerated degradation of another chimeric oncoprotein (EWS/FLI-1), possibly conferring therapeutic benefit in Ewing’s sarcoma^91^. While the exact mechanisms by which mTOR stabilizes *TFE3*-fusion proteins remain uncertain, potential hypotheses include increased lysosomal activation and/or accelerated autophagic flux downstream of mTOR inhibition, and will be clarified in further studies.

In summary, we describe a novel transgenic mouse model of renal PEComas induced by *SFPQ-TFE3* expression, the most common *TFE3* gene fusion seen in human PEComas^9,30^. More dramatic than *PRCC-TFE3*, *SFPQ-TFE3* expression is sufficient to rapidly drive lineage plasticity in renal tubular cells, evidenced by renal lineage factor downregulation, loss of cytokeratin expression and melanocytic/lysosomal marker upregulation. These data provide some of the first definitive evidence that PEComas may derive from an epithelial cell. Like other fusion-related models of tRCC, *SFPQ-TFE3*-driven renal tumorigenesis is accompanied by early activation of the mTORC1 pathway and we demonstrate for the first time that that mTOR activation itself positively feeds back to reinforce *TFE3*-fusion expression. Future work will further probe the mechanism of epithelial lineage plasticity induced by *TFE3*-fusions, and test whether mTOR kinase inhibition may have therapeutic efficacy in human fusion-driven tRCC and PEComas.

## Acknowledgements

This research was supported in part by the CDMRP TSCRP grant W81XWH-22-1-0264 (TLL), CDMRP KCRP grant W81XWH-20-1-0843 (TLL), CDMRP KCRP grant W81XWH2210377 (KA), and the NCI Cancer Center Support Grant 5P30CA006973-52. This work was supported at Johns Hopkins in part by Dahan Translocation Carcinoma Fund and Joey’s Wings Foundation. This work relied on expertise provide by the Polycystic Kidney Disease Research Resource Consortium, U54 DK126114. This research was supported by the Intramural Research Program of the NIH, National Cancer Institute, Center for Cancer Research (WML). This project has been funded in whole or in part with Federal funds from the National Cancer Institute, National Institutes of Health, under Contract No. HHSN261201500003I. (LSS). The content of this publication does not necessarily reflect the views or policies of the Department of Health and Human Services, nor does mention of trade names, commercial products, or organizations imply endorsement by the U.S. Government.

## Contributions

T.L.L. and K.A. conceived the study. K.A., A.A. and T.L.L. drafted the manuscript. K.A., A.A., J.W., S.N.A., T.V., K.F., L.O., H.B.L., M.K., M.B., Y.O., P.O., T.W., A.Z.R., L.S.S., W.M.L., P.A. and T.L.L. completed the data collection and analysis. All authors critically reviewed the manuscript and agreed to submit for publication.

## Methods

### Cell culture

UOK cell lines and HK2 cells with stable, doxycycline-inducible expression of TFE3 proteins were a kind gift of Dr. W. Marston Linehan (NCI). HEK293 cells with stable doxycycline-inducible expression of TFE3 proteins were generated using Flp Recombinase-mediated integration, using the Flp-In T-Rex Core Kit (K6500-01, Invitrogen). UOK cells were maintained in DMEM high glucose medium (#11995065, Gibco) with l-glutamine, 10% heat-inactivated FBS (#SH30071.03HI, Hyclone), 1X MEM (#11130051, Gibco) and 1% penicillin/ streptomycin at 37°C in 5% CO_2_. Inducible HEK cells were maintained in DMEM high glucose medium, 10% FBS (Tet-approved; A4736401, Gibco), Blasticidin S (15 µg/ml) Hygromycin B (150 µg/ml) and 1% penicillin/ streptomycin. Inducible HK-2 cells were maintained in Advanced DMEM/F-12 (12634010, Thermo), 1.5% FBS (Tet-approved) 1X GlutaMAX (35050061, Thermo), Puromycin (0.8 µg/ml), Blasticidin S (2 µg/ml) and 1% penicillin/ streptomycin. For experiments involving amino acid starvation, cells were rinsed in PBS and incubated in amino acid-free DMEM (#MBS6120661, MyBioSource) supplemented with 10% dialyzed serum (#26400044, Gibco) and glucose (A24940-01, Thermo) for 60-90 min. For amino acid addback, cells were stimulated with a 1X mix of essential (#11130036, Thermo Fisher Scientific) and non-essential amino acids (#11140035, Thermo Fisher Scientific) and 200 mM L-Glutamine (#25030081, Thermo Fisher Scientific), added directly to starved cells for 30 min.

### CRISPR-Cas9 genome editing for TFE3

We designed single-guide RNA (sgRNA) for 3 target sequences in the human *TFE3* gene (GGCGATTCAACATTAACGACAGG, GCGACGCTCAACTTTGGAGAGGG, TCGCCTGCGACGCTCAACTTTGG) and cloned these into the lentiCRISPR v2 vector (Addgene #52961, Watertown, MA, USA). Lentivirus was produced as previously described ^1^ and HK-2*/SFPQ-TFE3* cells were infected for 48 h, selected with puromycin (1 μg/mL) for 10 days, and colonies established.

### Plasmids, Lentiviral transfections and RNAi

Cells were transiently transfected using Lipofectamine 3000 (L3000008, Thermo; plasmid transfections) or Lipofectamine RNAiMAX (13778075, Thermo; siRNA transfections) according to the transfection guidelines. The following siRNAs were used: *RHEB* -siGENOME Human RHEB siRNA SMARTpool (Dharmacon; M-009692-02-0005), *RRAGC-*(Sigma; SASI_Hs01_00190647), *RRAGD-*(Sigma; SASI_Hs01_00057779), *PAX8* (SASI_Hs01_00211299).

### Antibodies and Reagents

<u>Primary antibodies:</u> **4E-BP1** (#9644 CST; 1:2000), **CTSK** (#ab19027 Abcam; 1:1000), **FLCN** (#3697 CST; 1:1000), **GAPDH** (#5174 CST; 1:1000), GATA3(#5852 CST; 1:500), **GPNMB** (#38313 CST; 1:1000), **GPNMB** (Mouse specific) (#90205 CST; 1:1000), **Histone-3** (#4499 CST; 1:4000), **HNF4A** (#3113 CST; 1:500), **Keratin 8** (#ab53280 Abcam; 1:1000), **LC3 A, B** (#12741 CST; 1:2000), **MelanA** (#ab210546 Abcam; 1:1000), **PAX2** (#9666 CST; 1:500), **PAX8** (#59019 CST; 1:500), **PMEL** (#ab137078 Abcam; 1:1000), **Phospho-p70 S6 Kinase (Thr389)** (#9205 CST; 1:1000), **p70 S6 Kinase** (#9202 CST; 1:1000), **Phospho-S6 Ribosomal Protein (Ser235/236)** (#4858 CST; 1:2000), **Phospho-4E BP1 (Ser65)** (#9451 CST; 1:2000), **Phospho-4E BP1 (Thr37/46)** (#2855 CST; 1:1000), **p-TFEB(S122)** (#86843 CST; 1:2000), **p-TFEB(S211)** (#37681 CST; 1:2000), **Pan-Keratin (Type1)** (#83957 CST; 1:1000), **RAB7** (#9367 CST; 1:1000), **RHEB** (#13879 CST; 1:1000), **RRAGC** (#9480 CST; 1:1000), **RRAGD** (#4470 CST; 1:1000), **Synaptophysin** (#5461 CST; 1:1000), **S6 Ribosomal Protein** (#2317 CST; 1:2000), **TFE3** (#14779 CST; 1:4000), **TFE3** (#ABE1400 Sigma; 1:4000), **TFEB** (#4240 CST; 1:2000), **WT1** (#83535 CST; 1:1000).

<u>Reagents:</u> Torin1 (#14379 CST), Doxycycline hyclate (D9891, Sigma), Cell lysis Buffer (9803, Cell Signaling), RIPA buffer (R0278, Sigma), MiniCollect Tube 0.5/0.8ml CAT Serum Sep Clot Activator (450533, Greiner), BrdU (#550891, BD Pharmingen)

### Animal Studies

Animal protocols were approved by the JHU Animal Care and Use Committee. The following strains were used: **1)** *SFPQ-TFE3^LSL^* mice expressing the *SFPQ-TFE3* fusion downstream of a *LoxP-Stop-LoxP (LSL)* cassette, were generated by Taconic Biosciences. **2)** Mice hemizygous for the *<u>Ksp</u>*<u>-Cre recombinase</u> knockin gene (Strain Number: 012237) (The Jackson Laboratory). **3)** Tamoxifen-inducible, *Pax8* Cre-ER^T2^ mice were a kind gift of Dr. Athena Matakidou (Cancer Research, UK). **4)** *PRCC-TFE3^LSL^* mice expressing the *PRCC-TFE3* fusion downstream of a *LoxP-Stop-LoxP (LSL)* cassette, were a kind gift of Dr. W. Marston Linehan (NCI). Genomic DNA was isolated from tail snips and genotyping performed using the following primers:

1) *<u>SFPQ-TFE3^LSL^</u>* - Transgene Forward: 5’-CTT-TAT-TAG-CCA-GAA-GTC-AGA-TGC-3’

Transgene Reverse: 5’-TGG-AGG-ACA-TTC-TGA-TGG-AGG-AG-3’

WT Forward: 5’-CAC TTG CTC TCC CAA AGT CGC TC-3’

WT Reverse: 5’-ATA CTC CGA GGC GGA TCA CAA-3’

2) *<u>Ksp</u>*<u>-Cre</u>-Transgene Forward: 5’-GCA GAT CTG GCT CTC CAA AG-3’

Transgene Reverse: 5’-AGG CAA ATT TTG GTG TAC GG-3’

3) *<u>Pax8</u>* <u>Cre-ER^T2^</u>-Transgene Forward: 5’-TGC CAC GAC CAA GTG ACA GCA ATG-3’

Transgene Reverse: 5’-ACC AGA GAC GGA AAT CCA TCG CTC-3’

4) *<u>PRCC-TFE3^LSL^</u>* - Transgene Forward: 5’-TTC CCC TCG TGA TCT GCA AC-3’

Transgene Reverse: 5’-CTG GAA AGA CCG CGA AGA GT-3’

WT Forward: 5’-TTC CCC TCG TGA TCT GCA AC-3’

WT Reverse: 5’-TCA TGG AAA TCT CCG AGG CG-3’

#### BUN and Creatinine measurements

were performed using mouse serum by IDEXX BioAnalytics.

#### BrdU labeling

To measure *in vivo* proliferation, mice were injected i.p. with a single dose of BrdU (#550891, BD Pharmingen), at a dose of 100 mg/kg body weight, 3 hours prior to sacrificing. BrdU incorporation was detected by immunohistochemistry of paraffin-embedded sections using an anti-BrdU monoclonal antibody (#5292 CST; 1:200).

### Histology and immunostaining

Mouse kidneys were fixed in 10% neutral buffered formalin (Sigma-Aldrich), embedded in paraffin, sectioned at 4 µm and used for H&E staining and immunohistochemistry. TFE3, GPNMB, Melan A, PMEL, Phospho-S6 Ribosomal Protein (Ser235/236), Phospho-4E BP1 (Thr37/46), TFEB, PAX8, PAX2, Pan-Keratin, CK8, Vimentin α-SMA, Synaptophysin, BrDU, Ki67, and Phospho-Histone 3 IHC on murine tissues was performed on the Ventana Discovery ULTRA (version v12.31) (Ventana/ Roche) using hand-applied antibodies at the following concentrations: **TFE3** (Invitrogen PA5-54909; 1:5000), **GPNMB** (#90205 CST; 1:100)**, Melan A** (#ab210546 Abcam; 1:500), **PMEL** (#ab137078 Abcam; 1:100), **Phospho-S6 Ribosomal Protein (Ser235/236)** (#4858 CST; 1:200), **Phospho-4E BP1 (Thr37/46)** (#2855 CST; 1:800), **TFEB** (#A303-673A, Bethyl; 1:1000), **PAX8** (#ab191870 Abcam; 1:100)**, PAX2** (#ab79389 Abcam; 1:500)**, Pan-Keratin** (#83957 CST; 1:25), **CK8** (#ab53280 Abcam; 1:100), **Vimentin** (#5741 CST; 1:200), ***α*-SMA** (#19245 CST; 1:200), **Synaptophysin** (#36406 CST; 1:50), **BrdU** (#5292 CST; 1:200), **Ki67** (#12202 CST; 1:100), **p-Histone H3** (#06-570 Sigma; 1:500).

#### Quantification of nuclear Pax8 and Pax2 in murine renal tumors

Immuno-stained slides were digitally scanned (Nanozoomer, Hamamatsu). Using *STK* and *PTK* kidney dual stained slides (GPNMB with Teal and PAX8 with DAB), we trained the HALO Tissue Classifier (IndicaLabs) to attribute classes (GPNMB-positive tubules or GPNMB-negative tubules) based on the stains and textures of the tissue and generated annotations accordingly. A CytoNuclear v2.0.9 algorithm (HALO®, Indica Labs) was run in each annotation to detect PAX8-negative and -positive tubular cells and further grade the positive cells into three intensities. The resulted H-score was used to compare the PAX8 intensity between the two classes. To validate the classifier, the algorithm was run in manually annotated areas and its results were correlated to the auto-detected annotations (r:0.97, p<0.000). The analysis algorithm was visually validated. PAX2 staining was visually assessed and showed the same pattern of positivity and intensity as PAX8.

#### LTL/ DBA Immunofluorescence

LTL/DBA staining was performed on paraffin-embedded kidney sections that were deparaffinized and rehydrated according to standard histological protocols. Sections were immersed in Target Retrieval Solution (Dako, S169984-2), heated in a steamer for 20 minutes and blocked using a blocking buffer containing 5% fetal bovine serum (FBS), 0.5% bovine serum albumin (BSA), and 0.1% Triton X-100 for 30 minutes. Rhodamine-labeled Dolichos biflorus agglutinin (DBA) and fluorescein-labeled Lotus tetragonolobus lectin (LTL) (Vector Laboratories, #RL-1032 and #FL-1321, respectively) at 1:200 dilution, were incubated overnight at 4°C. Subsequently, the nuclei were counterstained with DAPI, followed by three washes with PBST (PBS with Tween-20). Finally, the sections were mounted using Fluoromount-G™ Mounting Medium (Invitrogen, #00-4958-02). Images were captured using a Nikon W-1 spinning disk confocal microscope at the UMB-SOM Confocal Microscopy Core in Baltimore, Maryland.

### Cell and tissue lysates and immunoblotting

<u>Renal tumors:</u> were homogenized and lysed using the gentleMACS M Tubes/ Octo Dissociator in ice-cold RIPA lysis buffer. <u>Cells</u> were lysed in RIPA buffer or cell lysis buffer (for p-TFEB expression) (#9803, Cell Signaling). All lysis buffers were supplemented with NaVO_4_ (1 mM), NaF (1 mM) and 10 μl Halt Protease and Phosphatase Inhibitor Cocktail (#78440, Thermo Fisher Scientific). Lysates were centrifuged at 21,000 rpm for 10 minutes at 4°C and supernatants collected. Protein concentrations were quantified using the BCA Protein Assay Kit (#23225, Pierce), and protein was resolved on 4-12% Bis-Tris SDS-PAGE gel (Thermo Fisher Scientific). Protein was transferred to nitrocellulose membranes (Amersham Bioscience), blocked for 1h at room temperature in 5% nonfat milk in 1X TBS-T and then incubated overnight with a primary antibody diluted in 5% BSA or milk in 1X TBS-T. The secondary antibodies used were anti-rabbit or anti-mouse immunoglobulin as appropriate (Cell Signaling) and diluted at 1:1000 in 5% nonfat milk in 1X TBS-T. Blots were developed using a chemiluminescent development solution (Super Signal West Femto, Pierce) and bands were imaged on a chemiluminescent imaging system (ChemiDoc Touch imaging System using the ImageLab Touch Software (version 2.3.0.07) (Bio-Rad) or MicroChemi Chemiluminescent imager using the GelCapture Software (version 2.2.2.0) (FroggaBio Inc.). Digital images were quantified using Image J (version 1.52p) and all bands were normalized to their respective β-actin or GAPDH expression levels as loading controls.

<u>Nuclear lysates</u> were prepared using the PARIS kit (AM1921, Thermo Fisher Scientific) according to manufacturer’s instructions. Digital images were quantified using Image J and all bands were normalized to their respective Lamin, Histone H3 or Fibrillarin levels as loading controls. Statistical analysis was performed using Student’s unpaired t-test or one-way ANOVA.

### RNA isolation and quantitative real-time RT-PCR

Total cellular RNA was extracted using RNeasy Mini kit (#74104, Qiagen) according to manufacturer’s instructions. RNA was converted to cDNA using SuperScript III First-Strand Synthesis System (#18080051, Thermo Fisher Scientific) according to manufacturer’s instructions. mRNA levels were quantified using an ABI Prism 7900HT Real-time PCR system using the SDS (Sequence Detection System) software (version 2.4) (Applied Biosystems) with the following **<u>human</u>** primers and probes: **PAX2** (Hs01057416_m1), **PAX8 (**Hs00247586_m1), **HNF1B (**Hs01001602_m1), **LHX1 (**Hs00232144_m1), **GATA3 (**Hs00231122_m1), **PMEL (**Hs00173854_m1), **GPNMB (**Hs01095669_m1) and **GAPDH (**Hs02786624_g1) and **<u>mouse</u>** primers and probes: **RRAGC** (Mm00600306_m1), **RRAGD (**Mm00546741_m1), **GAPDH (**Mm99999915_g1). Threshold cycle (Ct) was obtained from the PCR reaction curves and mRNA levels were quantitated using the comparative Ct method with actin or Gapdh mRNA serving as the reference. Statistical analysis was performed using Student’s unpaired t-test.

### Immunocytochemistry

Cells seeded on fibronectin-coated coverslips were fixed in 4% PFA for 15 minutes at room temperature. Following three rinses in 1X PBS, cells were permeabilized and blocked in a buffer containing 1X PBS, 5% normal donkey serum and 0.3% Triton X-100. For immunofluorescence, coverslips were incubated with the indicated primary antibodies overnight at 4°C in antibody dilution buffer (ADB) containing 1X PBS, 1% BSA and 0.3% Triton X-100. After 3 rinses of 1X PBS, coverslips were incubated with secondary antibodies (Alexafluor-488 or Alexafluor-594 conjugated, anti-Rabbit or anti-Mouse IgG, Thermo Fisher Scientific) in ADB at a dilution of 1:200 for 1 hour at room temperature. Nuclei were counterstained with DAPI and coverslips visualized using an Olympus BX41 epifluorescence microscope using DP Controller software (version 3.2.1.276) (Olympus, Center Valley, PA).

### RNA sequencing and data analysis

RNA sequencing of triplicate cell line and xenograft replicates was performed at Novogene and carried out as previously described^2^. Raw RNAseq counts were Fragments Per Kilobase of transcript per Million mapped reads (FPKM)-normalized for data visualization in R (v4.3.2). Raw counts were also imputed in DESeq2 in R to determine differentially expressed genes. Log2 fold-changes, p-values, and adjusted p-values (false discovery rate method, FDR) were obtained for all genes and comparisons.

### Gene Set Enrichment Analysis (GSEA)

Raw counts were used as input for Gene Set Enrichment Analysis (GSEA, http://www.broad.mit.edu/gsea/). We employed the curated Hallmarks pathways (https://www.gsea-msigdb.org/gsea/msigdb/human/genesets.jsp?collection=H) to identify differential regulation of pathways in our comparison groups. In addition, curated sets from published studies were used to identify significantly enriched pathways in our study groups from data deposited in: a) dbGap under accession code phs001357.v1.p1.^3^, b) GE0 GSE252047^4^, or c) GEO GSE130072^5^. Negative NES indicated negatively enriched pathways in our comparison group vs. control. Q-value cutoffs were set to 0.1. Pan-Cancer-normalized RNAseq data from TCGA was downloaded from TCGA Pan-Cancer publication portal (https://gdc.cancer.gov/about-data/publications/panimmune). KIRP and KIRC RNAseq data were previously normalized with the methods Fragments Per Kilobase of transcript per Million mapped reads (FPKM) and FPKM Upper Quartile (FPKM-UQ) by the TCGA research team. Gene expression data was compared using Wilcoxon rank sum tests adjusted with multiple comparisons, where applicable. GSEA analyses were conducted with raw RNAseq counts using custom pathways and standard enrichment molecular signatures database. All analyses were performed in R v4.3.1.

### Statistics and Reproducibility

For RNA and protein quantification, statistical significance was determined using the unpaired, two-tailed Student’s t-test when comparing two experimental groups, or with one-way ANOVA with Dunnett’s or Bonferroni’s correction when comparing 3 or more experimental groups. Kaplan-Meier survival analyses were performed by Log-rank (Mantel-Cox) test. Statistical analyses of kidney to body weight ratios, BUN levels, Serum creatinine levels, BrdU incorporation and positivity, Ki67 positivity and Phosphorylated-Histone H3 (pH3) positivity were performed by Mann-Whitney test. Digital quantification of mean nuclear Pax8 and Pax2 H-scores in murine renal tumors was analyzed by two-tailed Mann-Whitney test. Gene expression data (FPKM plots) were analyzed by Wilcoxon rank sum test adjusted with multiple comparisons using the false discovery rate (FDR) method). Mean values were performed in GraphPad Prism (version 8.2.1). *p*-values of <0.05 were considered statistically significant. All experiments were repeated at least three times (independent biological replicates) with similar results. Additionally, all experiments were performed using multiple litters of mice and/or cellular replicates and using multiple orthogonal techniques to ensure rigor. For example: a) PAX2/PAX8 nuclear localization was confirmed by immunofluorescence (cells), IHC (mouse renal tumors) and immunoblotting of nuclear-cytoplasmic fractions.

## References

1 Kauffman, E. C. et al. Molecular genetics and cellular features of TFE3 and TFEB fusion kidney cancers. Nat Rev Urol 11, 465–475 (2014). 10.1038/nrurol.2014.162

2 Moch, H. et al. The 2022 World Health Organization Classification of Tumours of the Urinary System and Male Genital Organs-Part A: Renal, Penile, and Testicular Tumours. Eur Urol 82, 458–468 (2022). 10.1016/j.eururo.2022.06.016

3 Baba, M. et al. TFE3 Xp11.2 Translocation Renal Cell Carcinoma Mouse Model Reveals Novel Therapeutic Targets and Identifies GPNMB as a Diagnostic Marker for Human Disease. Mol Cancer Res 17, 1613–1626 (2019). 10.1158/1541-7786.MCR-18-1235

4 Salles, D. C. et al. GPNMB expression identifies TSC1/2/mTOR-associated and MiT family translocation-driven renal neoplasms. J Pathol 257, 158–171 (2022). 10.1002/path.5875

5 Wei, S., Testa, J. R. & Argani, P. A review of neoplasms with MITF/MiT family translocations. Histol Histopathol 37, 311–321 (2022). 10.14670/HH-18-426

6 Agaimy, A. et al. TFE3-rearranged nonmelanotic renal PEComa: a case series expanding their phenotypic and fusion landscape. Histopathology (2024). 10.1111/his.15304

7 Wang, X. T. et al. SFPQ/PSF-TFE3 renal cell carcinoma: a clinicopathologic study emphasizing extended morphology and reviewing the differences between SFPQ-TFE3 RCC and the corresponding mesenchymal neoplasm despite an identical gene fusion. Hum Pathol 63, 190–200 (2017). 10.1016/j.humpath.2017.02.022

8 Argani, P. et al. TFE3-Fusion Variant Analysis Defines Specific Clinicopathologic Associations Among Xp11 Translocation Cancers. Am J Surg Pathol 40, 723–737 (2016). 10.1097/PAS.0000000000000631

9 Rao, Q. et al. PSF/SFPQ is a very common gene fusion partner in TFE3 rearrangement-associated perivascular epithelioid cell tumors (PEComas) and melanotic Xp11 translocation renal cancers: clinicopathologic, immunohistochemical, and molecular characteristics suggesting classification as a distinct entity. Am J Surg Pathol 39, 1181–1196 (2015). 10.1097/pas.0000000000000502

10 Patel, S. A. et al. The renal lineage factor PAX8 controls oncogenic signalling in kidney cancer. Nature 606, 999–1006 (2022). 10.1038/s41586-022-04809-8

11 Bleu, M. et al. PAX8 activates metabolic genes via enhancer elements in Renal Cell Carcinoma. Nat Commun 10, 3739 (2019). 10.1038/s41467-019-11672-1

12 Siroky, B. J. et al. Evidence for pericyte origin of TSC-associated renal angiomyolipomas and implications for angiotensin receptor inhibition therapy. Am J Physiol Renal Physiol 307, F560–570 (2014). 10.1152/ajprenal.00569.2013

13 Fernandez-Flores, A. Evidence on the neural crest origin of PEComas. Rom J Morphol Embryol 52, 7–13 (2011).

14 Delaney, S. P., Julian, L. M. & Stanford, W. L. The neural crest lineage as a driver of disease heterogeneity in Tuberous Sclerosis Complex and Lymphangioleiomyomatosis. Front Cell Dev Biol 2, 69 (2014). 10.3389/fcell.2014.00069

15 Goncalves, A. F. et al. Evidence of renal angiomyolipoma neoplastic stem cells arising from renal epithelial cells. Nat Commun 8, 1466 (2017). 10.1038/s41467-017-01514-3

16 Yu, W. et al. Xp11.2 translocation renal neoplasm with features of TFE3 rearrangement associated renal cell carcinoma and Xp11 translocation renal mesenchymal tumor with melanocytic differentiation harboring NONO-TFE3 fusion gene. Pathol Res Pract 215, 152521 (2019). 10.1016/j.prp.2019.152521

17 Argani, P. et al. Xp11 translocation renal cell carcinoma (RCC): extended immunohistochemical profile emphasizing novel RCC markers. Am J Surg Pathol 34, 1295–1303 (2010). 10.1097/PAS.0b013e3181e8ce5b

18 Asrani, K. et al. An mTORC1-mediated negative feedback loop constrains amino acid-induced FLCN-Rag activation in renal cells with TSC2 loss. Nat Commun 13, 6808 (2022). 10.1038/s41467-022-34617-7

19 Alesi, N. et al. TFEB drives mTORC1 hyperactivation and kidney disease in Tuberous Sclerosis Complex. Nat Commun 15, 406 (2024). 10.1038/s41467-023-44229-4

20 Argani, P. et al. A distinctive subset of PEComas harbors TFE3 gene fusions. Am J Surg Pathol 34, 1395–1406 (2010). 10.1097/PAS.0b013e3181f17ac0

21 Malinowska, I. et al. Perivascular epithelioid cell tumors (PEComas) harboring TFE3 gene rearrangements lack the TSC2 alterations characteristic of conventional PEComas: further evidence for a biological distinction. Am J Surg Pathol 36, 783–784 (2012). 10.1097/PAS.0b013e31824a8a37

22 Akumalla, S. et al. Characterization of Clinical Cases of Malignant PEComa via Comprehensive Genomic Profiling of DNA and RNA. Oncology 98, 905–912 (2020). 10.1159/000510241

23 Sun, G. et al. Integrated exome and RNA sequencing of TFE3-translocation renal cell carcinoma. Nat Commun 12, 5262 (2021). 10.1038/s41467-021-25618-z

24 Qu, Y. et al. Proteogenomic characterization of MiT family translocation renal cell carcinoma. Nat Commun 13, 7494 (2022). 10.1038/s41467-022-34460-w

25 Prakasam, G. et al. Comparative genomics incorporating translocation renal cell carcinoma mouse model reveals molecular mechanisms of tumorigenesis. J Clin Invest 134 (2024). 10.1172/jci170559

26 Fang, R. et al. Nuclear translocation of ASPL-TFE3 fusion protein creates favorable metabolism by mediating autophagy in translocation renal cell carcinoma. Oncogene 40, 3303–3317 (2021). 10.1038/s41388-021-01776-8

27 Damayanti, N. P. et al. Therapeutic Targeting of TFE3/IRS-1/PI3K/mTOR Axis in Translocation Renal Cell Carcinoma. Clin Cancer Res 24, 5977–5989 (2018). 10.1158/1078-0432.CCR-18-0269

28 Napolitano, G. et al. A substrate-specific mTORC1 pathway underlies Birt-Hogg-Dube syndrome. Nature 585, 597–602 (2020). 10.1038/s41586-020-2444-0

29 Di Malta, C. et al. Transcriptional activation of RagD GTPase controls mTORC1 and promotes cancer growth. Science 356, 1188–1192 (2017). 10.1126/science.aag2553

30 Argani, P. et al. TFE3 -Rearranged PEComa/PEComa-like Neoplasms : Report of 25 New Cases Expanding the Clinicopathologic Spectrum and Highlighting its Association With Prior Exposure to Chemotherapy. Am J Surg Pathol 48, 777–789 (2024). 10.1097/PAS.0000000000002218

31 Argani, P. et al. Melanotic Xp11 translocation renal cancers: a distinctive neoplasm with overlapping features of PEComa, carcinoma, and melanoma. Am J Surg Pathol 33, 609–619 (2009). 10.1097/PAS.0b013e31818fbdff

32 Shao, X., Johnson, J. E., Richardson, J. A., Hiesberger, T. & Igarashi, P. A minimal Ksp-cadherin promoter linked to a green fluorescent protein reporter gene exhibits tissue-specific expression in the developing kidney and genitourinary tract. J Am Soc Nephrol 13, 1824–1836 (2002). 10.1097/01.asn.0000016443.50138.cd

33 Espana-Agusti, J. et al. Generation and Characterisation of a Pax8-CreERT2 Transgenic Line and a Slc22a6-CreERT2 Knock-In Line for Inducible and Specific Genetic Manipulation of Renal Tubular Epithelial Cells. PLoS One 11, e0148055 (2016). 10.1371/journal.pone.0148055

34 Cancer Genome Atlas Research, N., et al. Comprehensive Molecular Characterization of Papillary Renal-Cell Carcinoma. N Engl J Med 374, 135–145 (2016). 10.1056/NEJMoa1505917

35 Martin, K. R. et al. The genomic landscape of tuberous sclerosis complex. Nat Commun 8, 15816 (2017). 10.1038/ncomms15816

36 Subramanian, A. et al. Gene set enrichment analysis: a knowledge-based approach for interpreting genome-wide expression profiles. Proc Natl Acad Sci U S A 102, 15545–15550 (2005). 10.1073/pnas.0506580102

37 Liberzon, A. et al. The Molecular Signatures Database (MSigDB) hallmark gene set collection. Cell Syst 1, 417–425 (2015). 10.1016/j.cels.2015.12.004

38 Napolitano, G., Di Malta, C. & Ballabio, A. Non-canonical mTORC1 signaling at the lysosome. Trends Cell Biol 32, 920–931 (2022). 10.1016/j.tcb.2022.04.012

39 Martina, J. A. et al. The nutrient-responsive transcription factor TFE3 promotes autophagy, lysosomal biogenesis, and clearance of cellular debris. Sci Signal 7, ra9 (2014). 10.1126/scisignal.2004754

40 Glykofridis, I. E. et al. Loss of FLCN-FNIP1/2 induces a non-canonical interferon response in human renal tubular epithelial cells. Elife 10 (2021). 10.7554/eLife.61630

41 Horiguchi, H. et al. Dual functions of angiopoietin-like protein 2 signaling in tumor progression and anti-tumor immunity. Genes Dev 33, 1641–1656 (2019). 10.1101/gad.329417.119

42 Sancak, Y. et al. The Rag GTPases bind raptor and mediate amino acid signaling to mTORC1. Science 320, 1496–1501 (2008). 10.1126/science.1157535

43 Gollwitzer, P., Grutzmacher, N., Wilhelm, S., Kummel, D. & Demetriades, C. A Rag GTPase dimer code defines the regulation of mTORC1 by amino acids. Nat Cell Biol 24, 1394–1406 (2022). 10.1038/s41556-022-00976-y

44 Ansermet, C. et al. Renal tubular peroxisomes are dispensable for normal kidney function. JCI Insight 7 (2022). 10.1172/jci.insight.155836

45 Gu, X. et al. PBRM1 loss in kidney cancer unbalances the proximal tubule master transcription factor hub to repress proximal tubule differentiation. Cell Rep 36, 109747 (2021). 10.1016/j.celrep.2021.109747

46 Bakouny, Z. et al. Integrative clinical and molecular characterization of translocation renal cell carcinoma. Cell Rep 38, 110190 (2022). 10.1016/j.celrep.2021.110190

47 Calcagni, A. et al. Modelling TFE renal cell carcinoma in mice reveals a critical role of WNT signaling. Elife 5 (2016). 10.7554/eLife.17047

48 Chen, J. et al. Deficiency of FLCN in mouse kidney led to development of polycystic kidneys and renal neoplasia. PLoS One 3, e3581 (2008). 10.1371/journal.pone.0003581

49 Baba, M. et al. Kidney-targeted Birt-Hogg-Dube gene inactivation in a mouse model: Erk1/2 and Akt-mTOR activation, cell hyperproliferation, and polycystic kidneys. J Natl Cancer Inst 100, 140–154 (2008). 10.1093/jnci/djm288

50 Piontek, K., Menezes, L. F., Garcia-Gonzalez, M. A., Huso, D. L. & Germino, G. G. A critical developmental switch defines the kinetics of kidney cyst formation after loss of Pkd1. Nat Med 13, 1490–1495 (2007). 10.1038/nm1675

51 Zhang, H. et al. Loss of Tsc1/Tsc2 activates mTOR and disrupts PI3K-Akt signaling through downregulation of PDGFR. J Clin Invest 112, 1223–1233 (2003). 10.1172/jci17222

52 Ishiguro, N. & Yoshida, H. ASPL-TFE3 Oncoprotein Regulates Cell Cycle Progression and Induces Cellular Senescence by Up-Regulating p21. Neoplasia 18, 626–635 (2016). 10.1016/j.neo.2016.08.001

53 Pietrobon, A., Yockell-Lelievre, J., Flood, T. A. & Stanford, W. L. Renal organoid modeling of tuberous sclerosis complex reveals lesion features arise from diverse developmental processes. Cell Rep 40, 111048 (2022). 10.1016/j.celrep.2022.111048

54 Cho, J. H. et al. Notch transactivates Rheb to maintain the multipotency of TSC-null cells. Nat Commun 8, 1848 (2017). 10.1038/s41467-017-01845-1

55 Kwiatkowski, D. J. et al. A mouse model of TSC1 reveals sex-dependent lethality from liver hemangiomas, and up-regulation of p70S6 kinase activity in Tsc1 null cells. Human molecular genetics 11, 525–534 (2002).

56 Onda, H., Lueck, A., Marks, P. W., Warren, H. B. & Kwiatkowski, D. J. Tsc2(+/-) mice develop tumors in multiple sites that express gelsolin and are influenced by genetic background. J Clin Invest 104, 687–695 (1999). 10.1172/JCI7319

57 Liang, N. et al. Regulation of YAP by mTOR and autophagy reveals a therapeutic target of tuberous sclerosis complex. J Exp Med 211, 2249–2263 (2014). 10.1084/jem.20140341

58 Becherucci, F. & Romagnani, P. Angiomyolipoma: A link between stemness and tumorigenesis in the kidney. Nat Rev Nephrol 14, 215–216 (2018). 10.1038/nrneph.2018.16

59 Quintanal-Villalonga, Á. et al. Lineage plasticity in cancer: a shared pathway of therapeutic resistance. Nat Rev Clin Oncol 17, 360–371 (2020). 10.1038/s41571-020-0340-z

60 Betschinger, J. et al. Exit from pluripotency is gated by intracellular redistribution of the bHLH transcription factor Tfe3. Cell 153, 335–347 (2013). 10.1016/j.cell.2013.03.012

61 Robson, E. J., He, S. J. & Eccles, M. R. A PANorama of PAX genes in cancer and development. Nat Rev Cancer 6, 52–62 (2006). 10.1038/nrc1778

62 Patel, S. R., Ranghini, E. & Dressler, G. R. Mechanisms of gene activation and repression by Pax proteins in the developing kidney. Pediatr Nephrol 29, 589–595 (2014). 10.1007/s00467-013-2603-8

63 Narlis, M., Grote, D., Gaitan, Y., Boualia, S. K. & Bouchard, M. Pax2 and pax8 regulate branching morphogenesis and nephron differentiation in the developing kidney. J Am Soc Nephrol 18, 1121–1129 (2007). 10.1681/ASN.2006070739

64 Bouchard, M., Souabni, A., Mandler, M., Neubuser, A. & Busslinger, M. Nephric lineage specification by Pax2 and Pax8. Genes Dev 16, 2958–2970 (2002). 10.1101/gad.240102

65 Grote, D., Souabni, A., Busslinger, M. & Bouchard, M. Pax 2/8-regulated Gata 3 expression is necessary for morphogenesis and guidance of the nephric duct in the developing kidney. Development 133, 53–61 (2006). 10.1242/dev.02184

66 Tong, G. X. et al. Expression of PAX8 in normal and neoplastic renal tissues: an immunohistochemical study. Mod Pathol 22, 1218–1227 (2009). 10.1038/modpathol.2009.88

67 Cai, Q. et al. Pax2 expression occurs in renal medullary epithelial cells in vivo and in cell culture, is osmoregulated, and promotes osmotic tolerance. Proc Natl Acad Sci U S A 102, 503–508 (2005). 10.1073/pnas.0408840102

68 Beamish, J. A. et al. Pax protein depletion in proximal tubules triggers conserved mechanisms of resistance to acute ischemic kidney injury preventing transition to chronic kidney disease. Kidney Int 105, 312–327 (2024). 10.1016/j.kint.2023.10.022

69 Laszczyk, A. M. et al. Pax2 and Pax8 Proteins Regulate Urea Transporters and Aquaporins to Control Urine Concentration in the Adult Kidney. J Am Soc Nephrol 31, 1212–1225 (2020). 10.1681/ASN.2019090962

70 Laury, A. R. et al. A comprehensive analysis of PAX8 expression in human epithelial tumors. Am J Surg Pathol 35, 816–826 (2011). 10.1097/PAS.0b013e318216c112

71 Daniel, L. et al. Pax-2 expression in adult renal tumors. Hum Pathol 32, 282–287 (2001). 10.1053/hupa.2001.22753

72 Bradford, S. T. J. et al. Identification of Pax protein inhibitors that suppress target gene expression and cancer cell proliferation. Cell Chem Biol 29, 412–422.e414 (2022). 10.1016/j.chembiol.2021.11.003

73 Goea, L. et al. Hnf1b renal expression directed by a distal enhancer responsive to Pax8. Sci Rep 12, 19921 (2022). 10.1038/s41598-022-21171-x

74 Gresh, L. et al. A transcriptional network in polycystic kidney disease. EMBO J 23, 1657–1668 (2004). 10.1038/sj.emboj.7600160

75 Sun, M. et al. HNF1B Loss Exacerbates the Development of Chromophobe Renal Cell Carcinomas. Cancer Res 77, 5313–5326 (2017). 10.1158/0008-5472.CAN-17-0986

76 Naiman, N. et al. Repression of Interstitial Identity in Nephron Progenitor Cells by Pax2 Establishes the Nephron-Interstitium Boundary during Kidney Development. Dev Cell 41, 349–365.e343 (2017). 10.1016/j.devcel.2017.04.022

77 Ng-Blichfeldt, J. P., Stewart, B. J., Clatworthy, M. R., Williams, J. M. & Röper, K. Identification of a core transcriptional program driving the human renal mesenchymal-to-epithelial transition. Dev Cell 59, 595–612.e598 (2024). 10.1016/j.devcel.2024.01.011

78 Pena-Llopis, S. et al. Regulation of TFEB and V-ATPases by mTORC1. EMBO J 30, 3242–3258 (2011). 10.1038/emboj.2011.257

79 Menon, S. et al. Spatial control of the TSC complex integrates insulin and nutrient regulation of mTORC1 at the lysosome. Cell 156, 771–785 (2014). 10.1016/j.cell.2013.11.049

80 Sancak, Y. et al. Ragulator-Rag complex targets mTORC1 to the lysosomal surface and is necessary for its activation by amino acids. Cell 141, 290–303 (2010). 10.1016/j.cell.2010.02.024

81 Zoncu, R. et al. mTORC1 senses lysosomal amino acids through an inside-out mechanism that requires the vacuolar H(+)-ATPase. Science 334, 678–683 (2011). 10.1126/science.1207056

82 Gollwitzer, P., Grützmacher, N., Wilhelm, S., Kümmel, D. & Demetriades, C. A Rag GTPase dimer code defines the regulation of mTORC1 by amino acids. Nat Cell Biol 24, 1394–1406 (2022). 10.1038/s41556-022-00976-y

83 Schlingmann, K. P. et al. mTOR-Activating Mutations in RRAGD Are Causative for Kidney Tubulopathy and Cardiomyopathy. J Am Soc Nephrol 32, 2885–2899 (2021). 10.1681/asn.2021030333

84 Sambri, I. et al. RagD auto-activating mutations impair MiT/TFE activity in kidney tubulopathy and cardiomyopathy syndrome. Nat Commun 14, 2775 (2023). 10.1038/s41467-023-38428-2

85 Kauffman, E. C. et al. Preclinical efficacy of dual mTORC1/2 inhibitor AZD8055 in renal cell carcinoma harboring a TFE3 gene fusion. BMC Cancer 19, 917 (2019). 10.1186/s12885-019-6096-0

86 Parikh, J., Coleman, T., Messias, N. & Brown, J. Temsirolimus in the treatment of renal cell carcinoma associated with Xp11.2 translocation/TFE gene fusion proteins: a case report and review of literature. Rare Tumors 1, e53 (2009). 10.4081/rt.2009.e53

87 Malouf, G. G. et al. Targeted agents in metastatic Xp11 translocation/TFE3 gene fusion renal cell carcinoma (RCC): a report from the Juvenile RCC Network. Ann Oncol 21, 1834–1838 (2010). 10.1093/annonc/mdq029

88 Rua Fernandez, O. R., et al. Renal Cell Carcinoma Associated With Xp11.2 Translocation/TFE3 Gene-fusion: A Long Response to mammalian target of rapamycin (mTOR) Inhibitors. Urology 117, 41–43 (2018). 10.1016/j.urology.2018.03.032

89 Testa, S., Bui, N. Q. & Ganjoo, K. N. Systemic Treatments and Molecular Biomarkers for Perivascular Epithelioid Cell Tumors: A Single-institution Retrospective Analysis. Cancer Res Commun 3, 1212–1223 (2023). 10.1158/2767-9764.CRC-23-0139

90 Takla, M., Keshri, S. & Rubinsztein, D. C. The post-translational regulation of transcription factor EB (TFEB) in health and disease. EMBO Rep 24, e57574 (2023). 10.15252/embr.202357574

91 May, W. A. et al. The Ewing’s sarcoma EWS/FLI-1 fusion gene encodes a more potent transcriptional activator and is a more powerful transforming gene than FLI-1. Mol Cell Biol 13, 7393–7398 (1993). 10.1128/mcb.13.12.7393-7398.1993

## References

1 Walker, A. et al. Nrf2 signaling and autophagy are complementary in protecting breast cancer cells during glucose deprivation. Free Radic Biol Med 120, 407–413 (2018). 10.1016/j.freeradbiomed.2018.04.009

2 Asrani, K. et al. Reciprocal YAP1 loss and INSM1 expression in neuroendocrine prostate cancer. J Pathol (2021). 10.1002/path.5781

3 Martin, K. R. et al. The genomic landscape of tuberous sclerosis complex. Nat Commun 8, 15816 (2017). 10.1038/ncomms15816

4 Prakasam, G. et al. Comparative genomics incorporating translocation renal cell carcinoma mouse model reveals molecular mechanisms of tumorigenesis. J Clin Invest 134 (2024). 10.1172/jci170559

5 Baba, M. et al. TFE3 Xp11.2 Translocation Renal Cell Carcinoma Mouse Model Reveals Novel Therapeutic Targets and Identifies GPNMB as a Diagnostic Marker for Human Disease. Mol Cancer Res 17, 1613–1626 (2019). 10.1158/1541-7786.MCR-18-1235

