## Supplemental Figures for "*SFPQ-TFE3* gene fusion reciprocally regulates mTORC1 activity and induces lineage plasticity in a novel mouse model of renal tumorigenesis"

### Supplementary Figure 1

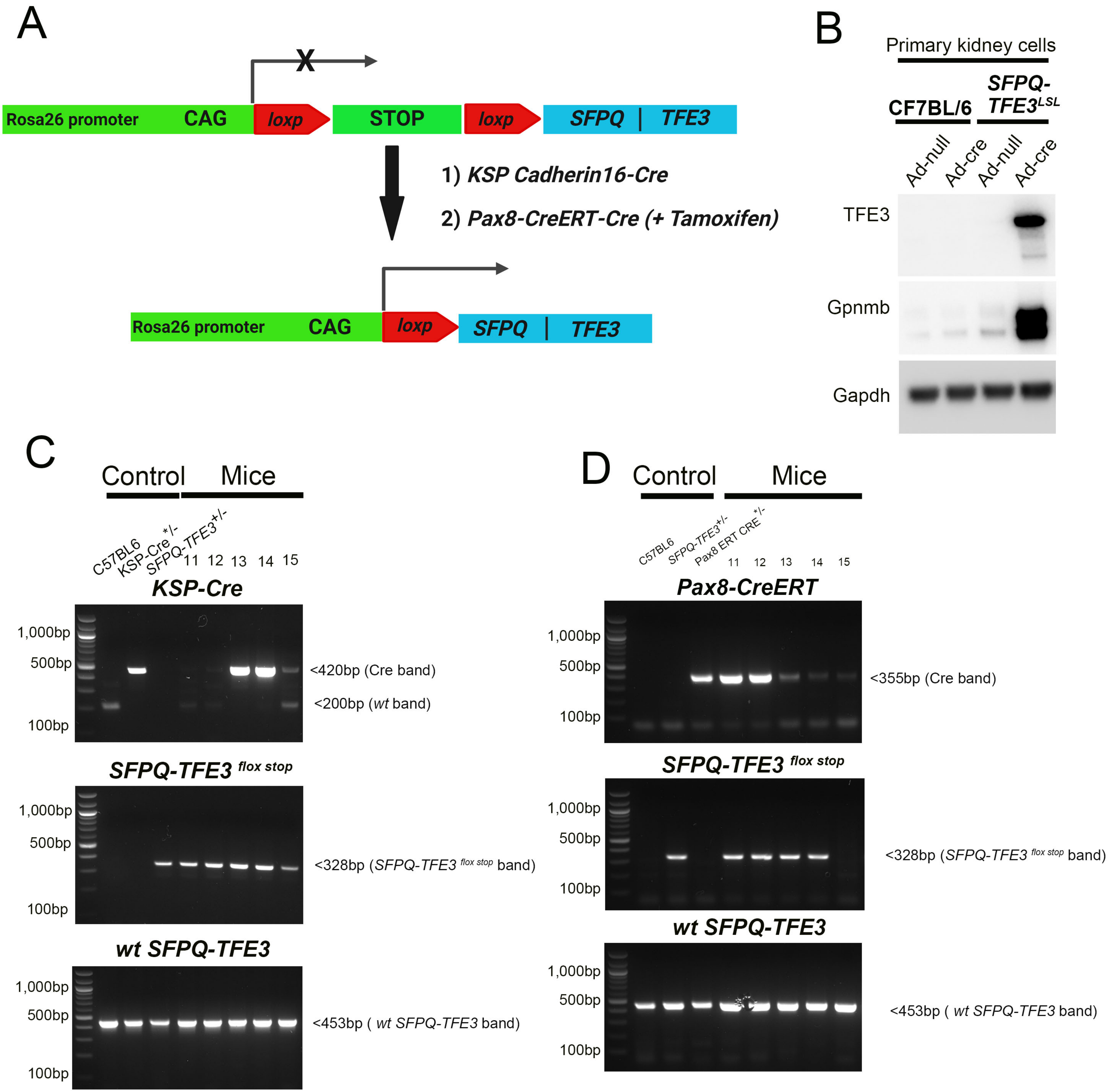

#### Supplemental Figure Legends

**Supplementary Figure S1:** ***Generation of an inducible murine allele of SFPQ-TFE3.*** **(A)** Schematic of the *SFPQ-TFE3* fusion transgene expression cassette. The human *SFPQ* CDS (exon 1-9) /human *TFE3* CDS (exon 5-10) (type 2 fusion), was cloned into intron 1 of the mouse *Rosa26* locus in reverse orientation, separated from the native CAG promoter by a stop sequence flanked by *LoxP* sites *(LSL).* Transgenic mouse expressing the human *SFPQ-TFE3* fusion protein, were crossed with either 1) Ksp-Cadherin 16 Cre mice or 2) Pax8 Cre-$ER^{T2}$ mice (with tamoxifen treatment), enabling conditional excision of the stop sequence upon Cre-mediated recombination and expression of the fusion transgene. **(B)** Immunoblotting of lysates from primary renal tubular epithelial cells from wild-type C57BL6 mice or *SFPQ-TFE3*$^{LSL}$ transgenic mice treated with control or Cre-recombinase expressing adenovirus *in vitro* for the indicated markers. Genotyping PCR from: **(C)** *SFPQ-TFE3*$^{LSL}$*; Ksp-Cre* mice or, **(D)** tamoxifen-treated, *SFPQ-TFE3 Pax8 ERT-Cre* (right panels), with expected molecular weights of PCR products indicated on right. Source data are provided as a Source data file.

### Supplementary Figure 2

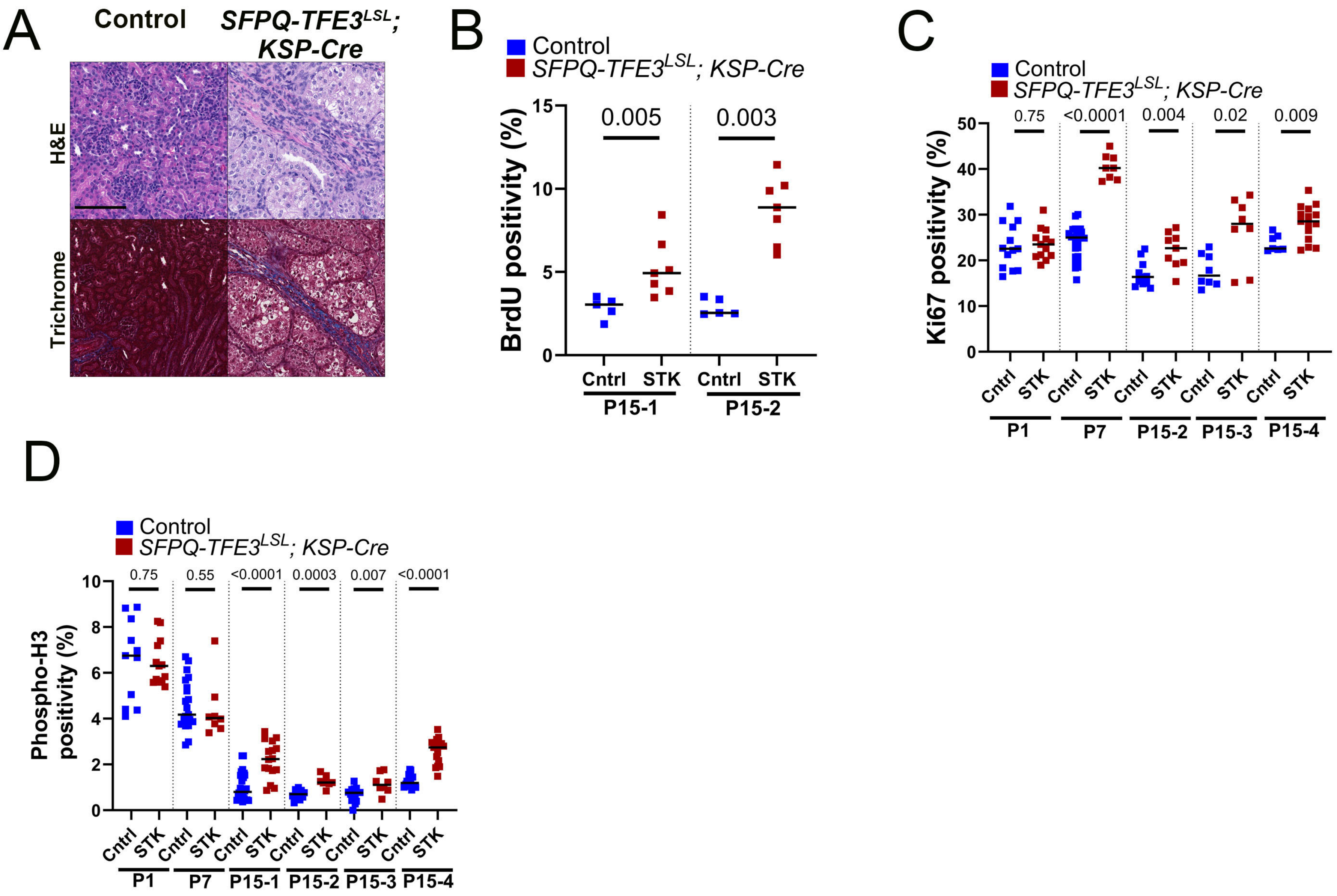

**Supplementary Figure S2:** ***Ksp-Cadherin Cre-mediated induction of SFPQ-TFE3 disrupts kidney development with renal failure and early neonatal death.*** **(A)** H&Es (top panels) and Masson's trichrome staining (bottom panels) showing features of inter-tubular fibrosis in *SFPQ-TFE3*$^{LSL}$*; Ksp-Cre* transgenic mice kidneys, compared to controls, at post-natal day 15. **(B)** BrdU incorporation and positivity in kidneys of age-matched, control and *SFPQ-TFE3*$^{LSL}$*; Ksp-Cre* transgenic mice at post-natal day 15, as measured by an IHC assay. The following numbers were analyzed in kidney sections from 2 pairs of mice: P15-1 [control=2, STK=2] and P15-2 [control=1, STK=1]. Data are presented as median. P values by Mann-Whitney test. **(C)** Ki67 positivity in kidneys of age-matched, control and *SFPQ-TFE3*$^{LSL}$*; Ksp-Cre* transgenic mice at post-natal days 1, 7 and 14, as measured by an IHC assay. P15 mice were analyzed in kidney sections from 3 separate mouse groups. The following numbers were analyzed: P1 [control=5, STK=6], P7 [control=2, STK=2], P15-2 [control=1, STK=1], P15-3 [control=2, STK=2], and P15-4 [control=3, STK=5]. Data are presented as median. P values by Mann-Whitney test. **(D)** Phosphorylated-Histone H3 (pH3) positivity in kidneys of age-matched, control and *SFPQ-TFE3*$^{LSL}$*; Ksp-Cre* transgenic mice at post-natal days 1, 7 and 14, as measured by an IHC assay. P15 mice were analyzed in kidney sections from 4 separate mouse groups. The following numbers were analyzed: P1 [control=5, STK=6], P7 [control=2, STK=2], P15-1 [control=2, STK=2], P15-2 [control=1, STK=1], P15-3 [control=2, STK=2], and P15-4 [control=3, STK=5]. Data are presented as median. P values by Mann-Whitney test. Source data are provided as a Source data file.

### Supplementary Figure 3

A B

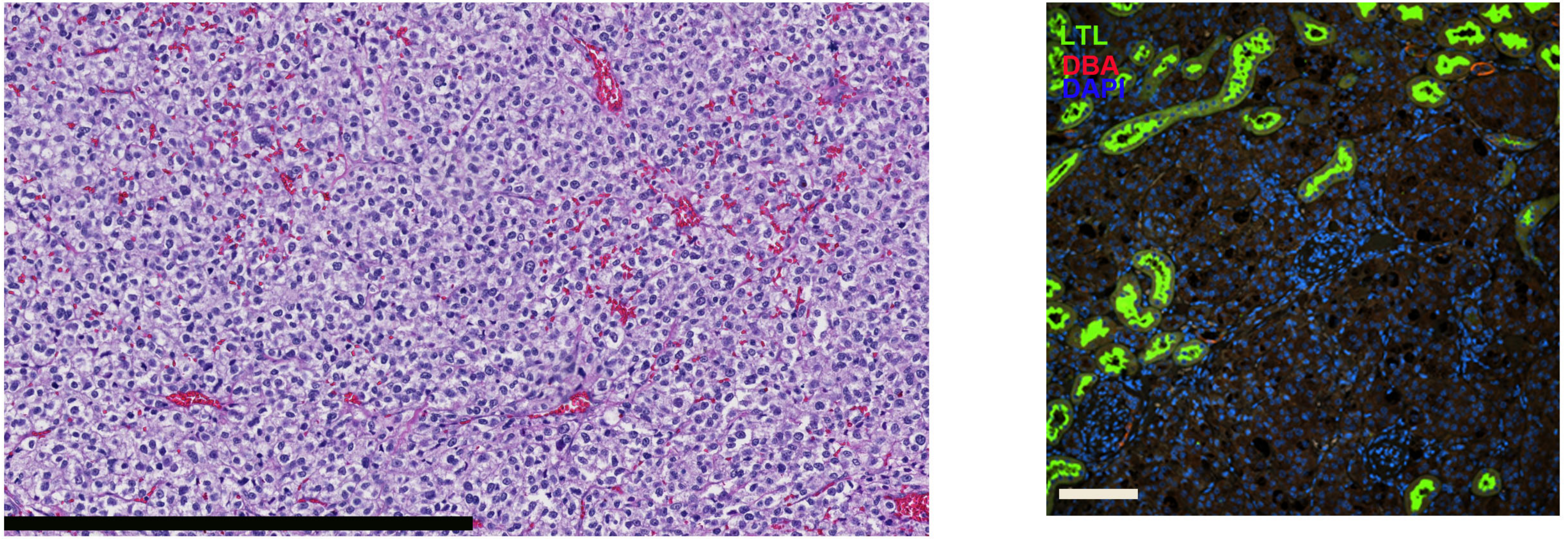

**Supplementary Figure S3:** ***Conditional post-natal, doxycycline-mediated induction of SFPQ-TFE3 in Pax8 Cre-ER$^{T2}$ mice induces renal tumor development.*** **(A)** H&E of a human *TFE3*-rearranged PEComa with nests of monomorphic epithelioid cells with clear to eosinophilic cytoplasm and large round nuclei. **(B)** Indirect immunofluorescence for LTL (green) and DBA (red) in *SFPQ-TFE3$^{LSL}$; Pax8-CreERT* transgenic mice, sacrificed at 3.5 months following injection of tamoxifen. Scale bar=100 µm. Source data are provided as a Source data file.

### Supplementary Figure 4

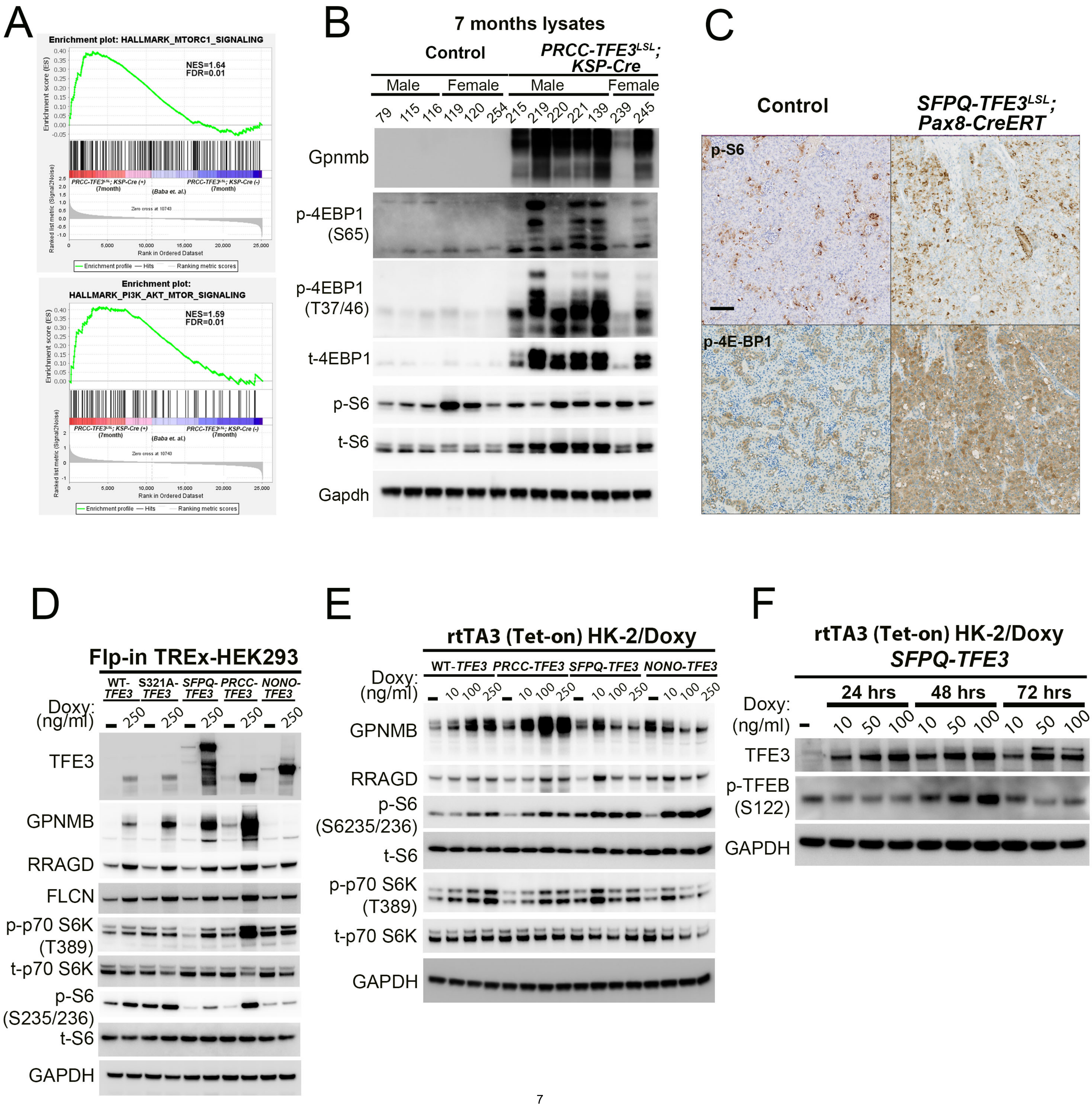

**Supplementary Figure S4:** ***mTORC1 signaling is activated in murine and human models of SFPQ-TFE3 fusion-RCC.*** **(A)** Gene Set Enrichment Analysis (GSEA) using Hallmark gene sets for MTORC1 signaling (upper panel) and PI3K/AKT/MTOR signaling (lower panel), in *PRCC-TFE3; KSP-Cre (+)* transgenic mice at 7 months, compared to controls[3]. **(B)** Immunoblotting of kidney lysates from control and *PRCC-TFE3; KSP-Cre (+)* transgenic mice at 7 months, for phosphorylation of mTORC1 substrates (p-4E-BP1[S65], p-4E-BP1[T37/46] and p-S6[S235/236]). **(C)** Representative immunohistochemistry (IHC) for p-S6[S235/236] (top row) and p-4E-BP1[T37/46] (bottom row) from age-matched, control and *SFPQ-TFE3*$^{LSL}$*; Pax8-CreERT* transgenic mice, sacrificed at 3.5 months following injection of tamoxifen. Scale bar= 100 μm. **(D)** Immunoblotting of lysates from HEK293 cells with doxycycline-inducible expression of *WT-TFE3*, *S321A-TFE3*, *SFPQ-TFE3, PRCC-TFE3* and *NONO-TFE3,* using the Flp-In-T-Rex™ system, for the indicated antibodies. Cells were untreated, or treated with the indicated doses of doxycycline for 48 hrs, prior to lysis and immunoblotting. **(E)** Immunoblotting of lysates from HK2 proximal tubular epithelial cells, with doxycycline-inducible expression of *WT-TFE3* and *TFE3* fusion proteins using the rtTA3 (Tet-on) system, for the indicated antibodies. Cells were untreated, or treated with the indicated doses of doxycycline for 72 hrs, prior to lysis and immunoblotting. **(F)** Immunoblotting of lysates from HK2/*SFPQ-TFE3* cells, treated with the indicated doses of doxycycline for 24, 48 or 72 hrs for expression of p-TFEB. Source data are provided as a Source data file.

### Supplementary Figure 5

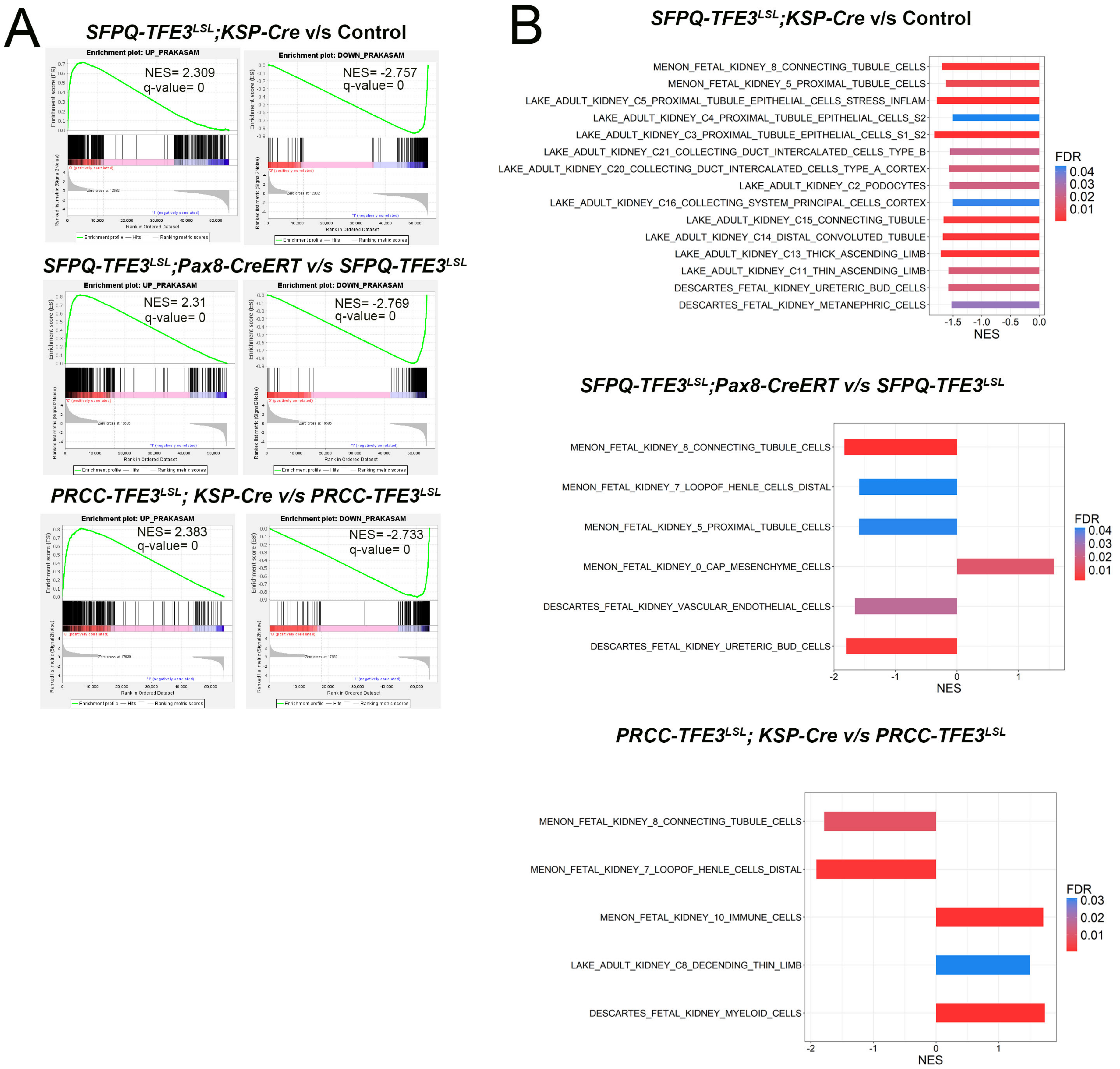

**Supplementary Figure S5:** ***Induction of SFPQ-TFE3 expression in human and murine renal tubular epithelial cells results in lineage plasticity with silencing of nephric lineage factors PAX8 and PAX2.*** **(A)** Gene Set Enrichment Analysis (GSEA) comparing 15-day *STK* (top panels)*,* 3.5-month tamoxifen-treated, *STP* (middle panels) and 7-month *PTK* (bottom panels), transgenic kidneys and their controls, for the core set of differentially expressed genes overlapping between human tRCC and the *ASPSCR1-TFE3* mouse model[25]. **(B)** Gene Set Enrichment Analysis (GSEA) comparing 15-day *STK* (top panels)*,* 3.5-month tamoxifen-treated, *STP* (middle panels) and 7-month *PTK* (bottom panels), transgenic kidneys and their controls, showing negative enrichment of genes associated with renal epithelial cell subsets from the cell type signature gene sets (C8).

### Supplementary Figure 6

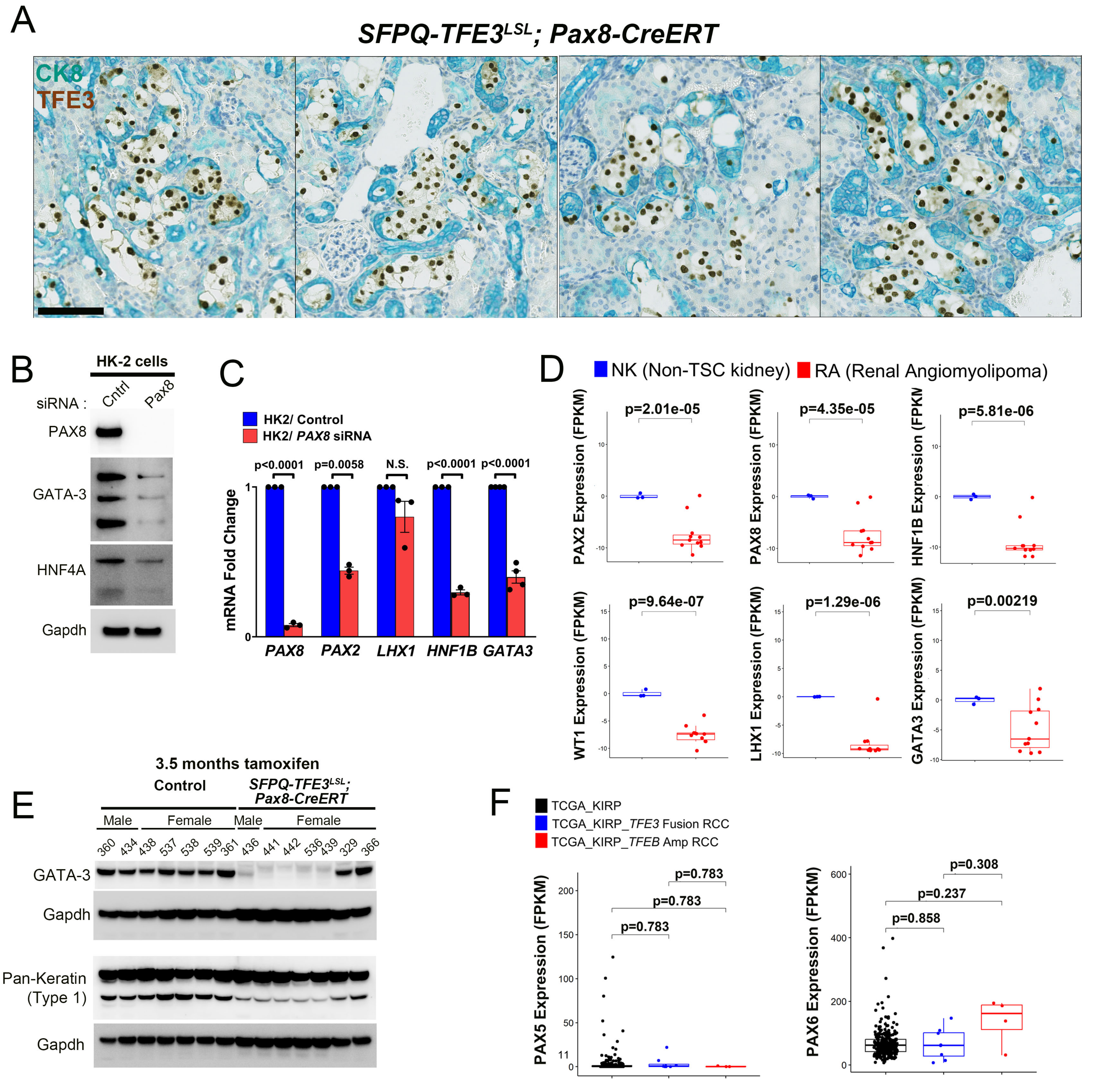

**Supplementary Figure S6:** ***Induction of SFPQ-TFE3 expression in human and murine renal tubular epithelial cells results in lineage plasticity with silencing of nephric lineage factors PAX8 and PAX2.*** **(A)** Dual IHC for TFE3 (brown) and CK8 (teal) in tamoxifen-injected, *SFPQ-TFE3*$^{LSL}$*; Pax8-CreERT* transgenic mice and age-matched, littermate controls, at 2 weeks following injection of tamoxifen. Scale bar = 100 µm. **(B)** Immunoblotting for GATA-3 and HNF4A in HK2 proximal tubular epithelial cells following treatment with control or *PAX8* siRNA for 48 hrs. **(C)** Quantitative real time PCR (qRT-PCR) for *PAX8, PAX2, HNF1B, LHX1* and *GATA3* in HK2 proximal tubular epithelial cells following treatment with control or *PAX8* siRNA for 48 hrs. (n=/>3, error bars represent SEM; p values by Student's T-test). **(D)** Comparison of *PAX2, PAX8, HNF1B, WT1, LHX1* and *GATA3* expression (FPKM) in RNA seq data from a panel of non-TSC normal kidneys (n=3) and renal angiomyolipomas with *TSC1/2* biallelic loss (n=11) [35]. (*P*-values indicated are by Wilcoxon rank sum test). **(E)** Immunoblotting of kidney lysates from tamoxifen-injected, control and *SFPQ-TFE3*$^{LSL}$*; Pax8-CreERT* transgenic mice at 3.5 months following injection of tamoxifen, for GATA-3. **(F)** Comparison of *PAX5* and *PAX6* gene expression in *TFE3* fusion-RCC (n=8) and *TFEB*-amplified RCC (n=4) to the remainder of papillary RCC cases without underlying *TFE3* fusions or *TFEB* amplifications (n=273) in the papillary RCC (KIRP) cohort from TCGA. Data are represented as a box-and-whisker plot. (*P*-values indicated are by Wilcoxon rank sum test adjusted with multiple comparisons using the false discovery rate (FDR) method). Source data are provided as a Source data file.

### Supplementary Figure 7

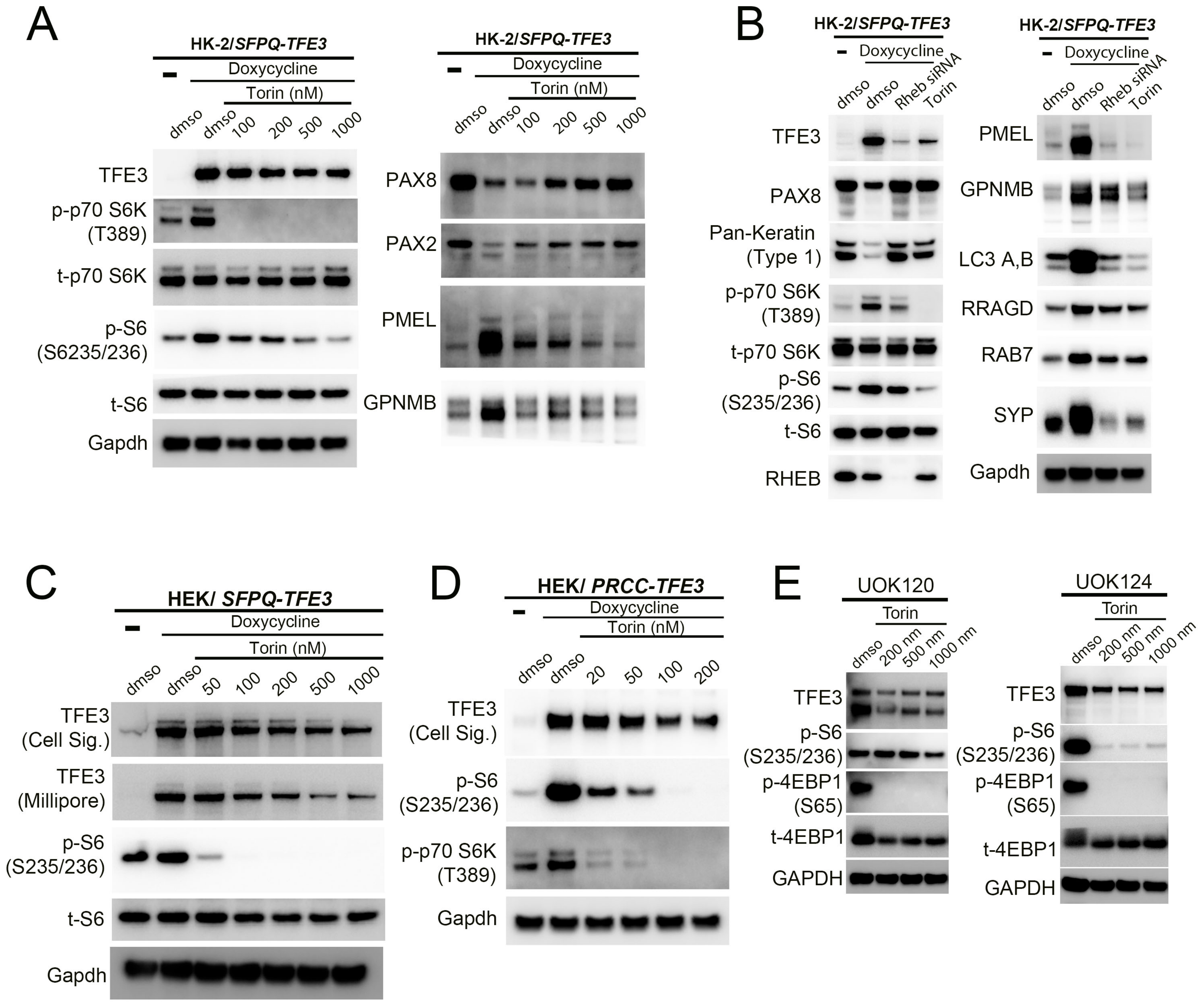

**Supplementary Figure S7:** ***mTORC1 inhibition rescues PAX2 and PAX8 expression and activation in human vitro models of SFPQ-TFE3 expression.*** **(A)** Immunoblotting of HK2 cells with doxycycline-inducible expression of *SFPQ-TFE3,* following treatment with the indicated doses of the mTOR kinase inhibitor torin1, for the indicated antibodies. (**B)** Immunoblotting of HK2 cells with doxycycline-inducible expression of *SFPQ-TFE3,* following treatment with *Rheb* siRNA or the mTOR kinase inhibitor torin1, for the indicated antibodies. **(C)** Immunoblotting of lysates from HEK293 cells with doxycycline-inducible expression of *SFPQ-TFE3,* following treatment with doses of the mTOR inhibitor torin1, for the indicated antibodies. **(D)** Immunoblotting of lysates from HEK293 cells with doxycycline-inducible expression of *PRCC-TFE3,* following treatment with doses of the mTOR inhibitor torin1, for the indicated antibodies. **(E)** Immunoblotting of lysates from *PRCC-TFE3* cell lines (UOK120 and124), following treatment with doses of the mTOR inhibitor torin1, for the indicated antibodies. Source data are provided as a Source data file.
